# Delineating single subject oscillatory brain networks with Spatio-Spectral Eigenmodes

**DOI:** 10.1101/2020.06.21.157412

**Authors:** Andrew J Quinn, Gary GR Green, Mark Hymers

**Affiliations:** Oxford Centre for Human Brain Activity, Wellcome Centre for Integrative Neuroimaging, University Department of Psychiatry, Warneford Hospital, Oxford, UK. OX3 7JX; York Neuroimaging Centre, The Biocentre, York Science Park, Heslington, York, UK. YO10 5NY; Department of Psychology, University of York, Heslington, York, UK. YO10 5DD

## Abstract

The spatial and spectral structure of oscillatory networks in the brain provide a readout of the underlying neuronal function. Within and between subject variability in these networks can be highly informative but also poses a considerable analytic challenge. Here, we describe a method that simultaneously estimate spectral and spatial network structure without assumptions about either feature distorting estimation of the other. This enables analyses exploring how variability in the frequency and spatial structure of oscillatory networks might vary both across the brain and across individuals. The method performs a modal decomposition of an autoregressive model to describe the oscillatory signals present within a time-series based on their peak frequency and damping time. Moreover, an alternate mathematical formulation for the system transfer function can be written in terms of these oscillatory modes; describing the spatial topography and network structure of each component. We define a set of Spatio-Spectral Eigenmodes (SSEs) from these parameters to provide a parsimonious description of oscillatory networks. Crucially, the SSEs preserve the rich between-subject variability and are constructed without pre-averaging within specified frequency bands or limiting analyses to single channels or regions. After validating the method on simulated data, we explore the structure of whole brain oscillatory networks in eyes-open resting state MEG data from the Human Connectome Project. We are able to show a wide variability in peak frequency and network structure of alpha oscillations and reveal a distinction between occipital ‘high-frequency alpha’ and parietal ‘low-frequency alpha’. The frequency difference between occipital and parietal alpha components is present within individual participants but is partially masked by larger between subject variability; a 10Hz oscillation may represent the high-frequency occipital component in one participant and the low-frequency parietal component in another. This rich characterisation of individual neural phenotypes has the potential to enhance analyses into the relationship between neural dynamics and a person’s behavioural, cognitive or clinical state

## 1 Introduction

The synchronised activity of neuronal populations is observable via the wide variety of oscillatory phenomena in electrophysiological recordings of brain function. Such oscillations are thought to reflect coordinated activity in neuronal networks carrying out specific functionality within the brain [1, 2]. These oscillatory signatures have a rich frequency spectrum that is often simplified, in analysis or interpretation, to a set of discrete band-limited components (typically delta, theta, alpha, beta and gamma). Whilst there is a strong basis for the functional, spatial and spectral separation of these components, analyses assuming a strong separation into *a-priori* defined bands risk overlooking spatial and spectral variability both within and between subjects.

For example, the alpha oscillation is often characterised as an 7–13Hz signal originating from occipital cortex [3, 4] whose function has been associated with a wide range of cognitive and clinical states [5, 6]. Yet there is strong and growing evidence that alpha oscillations are not homogeneous in across different frequencies, brain regions or individual participants. The lower and higher edges of the 7–13Hz alpha band have distinct task responses indicating that they shown to relate to different aspects of cognition [5, 7]. Individual Alpha Frequency (IAF) is variable across populations [7] and modulated by task demands within individuals [8]. Moreover, IAF may be a valuable clinical marker; the slowing of alpha peak frequency is a robust characteristic of both Alzheimer’s Disease and Mild Cognitive Impairment [9–15]. The spatial origin of the alpha rhythm is typically localised to the midline occipito-parietal and occipital cortex [16, 17] although individual subjects’ alpha networks show a wide variability in topography [18]. This network variability is heritable [19] and likely to reflect biologically relevant subject differences. Some of this spatial variability may arise from functionally distinct alpha generators in different brain regions [5, 20]. For instance, distinctions have been shown between occipito-parietal and occipito-temporal alpha [21], as well as alpha arising from visual and parietal sources [20].

These lines of evidence emphasise the functional relevance of spatial and spectral variability in neuronal oscillations whilst illustrating the difficulty of untangling the many sources of within and between subject variability. It remains a substantial analytic challenge to characterise variability in both the peak frequency, spatial distribution and network structure of neuronal rhythms. We present a novel, data-driven approach which characterises both the spatial and spectral structure of an oscillatory network. We use the modal decomposition [22, 23] of a multivariate autoregressive (MVAR) model to define a set of Spatio-Spectral Eigenmodes (SSEs). Each mode contains a unimodal (single peak) frequency response whose dynamical importance is represented by a damping time; rapidly damped modes will be quickly extinguished and contribute less to the observed dynamics in the data. Here, we show that the contribution of each mode to the system transfer function can be computed from the parameters of each mode. Both the peak oscillatory frequency and spatial representation are simultaneously estimated within each SSE, without needing to impose arbitrary *a priori* frequency bands or spatial regions of interest. Each SSE is a property of the whole system with a contribution to the whole network and whole power spectrum.

In this paper, we apply the Spatio-Spectral Eigenmodes to explore macro-scale spatial and spectral variability in oscillatory resting state networks both within and between subjects. The MVAR modal decomposition allows us to explore oscillatory network structure without distorting the peak frequency of each individual subject. The method is first validated with simulations before being applied to resting state MEG data from the Human Connectome Project [24, 25]. Source time-courses are estimated from the pre-processed sensor data using a LCMV-beamformer [26] before voxels are combined within regions of the Automated Anatomical Labelling (AAL) atlas [27]. This source-parcellated data is described with an MVAR before the modal decomposition is used to describe the oscillatory features in the data. We estimate the source power distribution from the MVAR parameters and identify dynamically important modes based on the damping times. We describe the principal components of spatial variability across the dynamically important SSEs revealing large-scale patterns in network structure across the whole brain. Importantly, though the individual variability in peak frequency is of a similar magnitude to the spatial variability, this method is able to show that the frequency difference within different gradients are largely consistent within individuals, even though the overall IAF is highly variable between individuals.

## 2 Results

### 2.1 Spatio-Spectral Eigenmodes from Multivariate Autoregressive models

Here, we give an overview of a standard approach to Multivariate Autoregressive (MVAR) modelling and spectrum estimation before outlining the the modal decomposition and definition of Spatio-Spectral Eigenmodes. An illustrative summary of the analysis pipeline used is given in figure 3

#### 2.1.1 Spectrum estimation from Multivariate Autoregressive Models

We start with a vector time series, **x**(t) with *m* channels *x*_1_(*t*)*, x*_2_(*t*)*, …, x_m_*(*t*), *t* ∈ 1, 2*, … , T*. Time-lagged linear dependencies within and between the channels can be characterised with an MVAR model of order *p*.

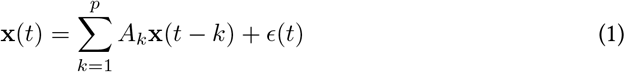

where *A_k_* is an *m × m* array of regression parameters at lag *k* and *ϵ* is an *m*-variate white noise process. This is a form of linear time-invariant (LTI) system in which future values of **x**(*t*) are predicted from a linearly weighed combination of its past values. The parameter matrix *A_k_* contains these linear dependencies between the past and future values of the time-series at a given lag, *k*. The off-diagonal elements of *A_k_* describe the degree to which the different channels within the system contain lagged interactions. *A* (without subscript) denotes the 3-dimensional parameter matrix containing *A_k_* for all fitted values of *k* (from 1 to *p*).

The interactions described by *A* may be expressed in the frequency domain by computing the system transfer function *H* as a function of frequency *f*. The transfer function describes the ratio of the input to a system to the output of the system and is computed from the *z*-transform of the *A* matrix.

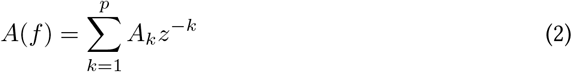

where

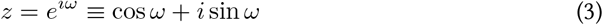

and

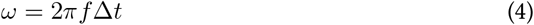

Δ*t* denotes the sampling interval, *ω* the normalised frequency in radians and *f* the frequency in Hertz. Equation 2 can be evaluated for any value of *z* in the complex plane. Here, we only evaluate *z* on the unit circle (where |*z*| = 1) as the output of these points can be directly related to an oscillatory frequency *f* and equation 2 is equivalent to the discrete time Fourier transform.

The power spectrum of **x**(*t*) can be computed from the frequency transform of the autoregressive parameters *A* via the transfer function *H*(*f*) and the residual covariance matrix Σ.

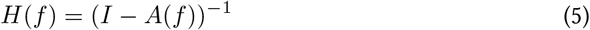

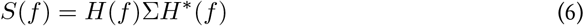

Where *H*^∗^(*f*) denotes the complex conjugate transform of *H*(*f*). *S*(*f*) contains the power spectrum of each of the *m* regions in the diagonal and the cross-spectrum between regions in the off-diagonal terms. The properties of *A*(*f*), *H*(*f*) and *S*(*f*) form the basis of a range of spectral connectivity metrics including Magnitude Squared Coherence, Geweke-Granger causality, Directed Transfer Function and Partial Directed Coherence [28, 29]. In practise, analyses will typically compute these metrics across a range of frequencies before integrating between specified frequency bands to isolate frequency specific structure.

#### 2.1.2 MVAR Modal Decomposition

An autoregressive model is a form of Infinite Impulse Response (IIR) filter whose spectral characteristics are completely described by the polynomial roots of its parameters. These roots directly relate to resonances in *H* and describe how the filter extracts an input at frequency *f* to obtain the filter output. This is well characterised for univariate systems and can be generalised to multivariate systems to provide an intuitive description of the frequency information contained in an MVAR model. This modal representation of the transfer function can then be used to simultaneously [23] explore the peak frequency and spatial structure of brain networks. The modal decomposition of MVAR coefficients is closely related to linear filter theory.

To perform the modal decomposition, we first rewrite the order-*p A* matrix as an order 1 system in a square block form. The autoregressive model in equation 1 can be restructured into a blocked form using a delay embedding of *X*(*t*) = **x**(*t*), **x**(*t*1)*, …*, **x**(*t − p*) and the companion form *C* of the MVAR parameter matrix [23].

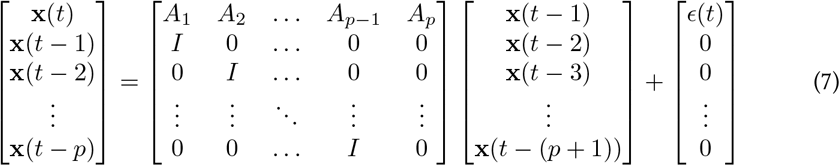

*C* is a blocked *mp × mp* matrix with the sparse *p* − 1 rows at the bottom shifting the corresponding rows in *X*(*t* − 1) down to create space for the *x*(*t*) in the prediction. The simplified matrix form of this equation

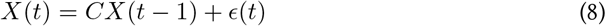

is of almost identical form to an order 1 autoregressive model in the standard formulation in equation 1. The eigendecomposition of the square parameter matrix *C* then yields *λ*, *V* and *W* as the eigenvalues, right eigenvectors and left eigenvectors respectively. The eigenvalues *λ* are the roots of the characteristic equation of the matrix *C* and as such directly define the frequency response of the pole. The characteristic frequency of each pole can be calculated as:

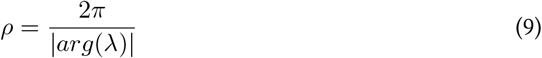

Oscillations are represented by complex conjugate pairs of poles within *λ* whilst single poles lying on the real line represent non-oscillatory parts of the signal. The damping time of a mode is also computed from its eigenvalue:

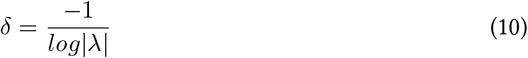

This describes the rate at which the amplitude of an oscillation would drop to zero if the system were energised with an impulse response. Longer damping times indicate less damped modes which will oscillate for longer durations following a single input. Short damping times indicate that the behaviour of the mode is quickly extinguished once the system is energised.

The complex valued eigenvector matrices *W* and *V* are the same *mp × mp* size as *C*. They have a specific Vandermonde structure in which a row contains *p* blocks of *m* values raised to successive powers of their corresponding eigenvalue *λ_j_*.

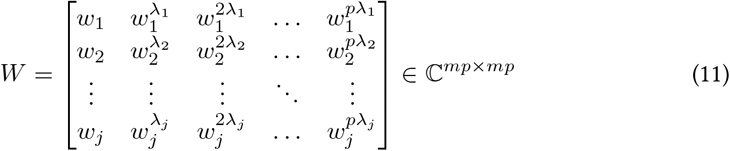

Due to the repeating structure in rows of *W*, we reduce analysis of the eigenvector of a mode to a vector of the first *m* values in each row (*w_j_*(1 : *m*) or *v_j_*(1 : *m*)). This reduced vector (also called mode shape) describes the structure of the resonance across the *m* dimensions of the input.

#### 2.1.3 The modal form of the transfer function

When using autoregressive models for spectrum analysis, the transfer function is typically estimated from the Fourier transform of the time-domain parameters *A* (equation 2). Here, we show that it may equivalently be computed from the modes from the eigenvalue decomposition. The eigenvalues and eigenvectors defined above form the parameters of a partial fraction expansion of the transfer function. This converts the transfer function from a ratio of two long polynomials to the sum across a set of fractions with simple denominators. A modal form of the transfer function can then be defined as a summation of a ratio of the properties of the *mp* modes (a full derivation is included in appendix 4.1).

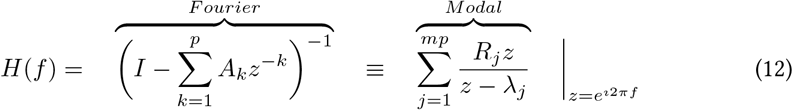

where *λ* is the modal eigenvalue and *R_j_* is the mode residue matrix. The Fourier form comes from substituting equation 2 into equation 5. In the modal form, the mode residue *R* is the coefficient of each term in the expansion (and distinct from the residuals of the autoregressive model fit) computed from the outer product of the first *m* terms in the left and right eigenvectors 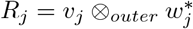 where ^∗^ denotes the complex conjugate. *R_j_* is then an *m* × *m* matrix whose elements are coefficients denoting the strength of the mode at each node and connection in the system. In other words, it acts to project the oscillation defined by *λ_j_* in the signal within each node and the connections between them. When all *mp* modes are included in the summation, the Fourier and Modal forms of *H* are exactly equivalent. The modal form is related to Gilbert’s Realisation [30, 31] which expresses a rational transfer function as a partial fraction expansion.

This modal form of *H* has several benefits. Firstly, the relation to the other modal parameters provides important context to *H*. Though we can evaluate *H* at any frequency up to the Nyquist limit, the resolution of the power spectrum is limited by the number of modes. A decomposition with a higher model order will have more modes in its decomposition and therefore a richer spectral structure. Secondly, as the modal form is a linear summation across modes, the contribution of a single resonance can be easily isolated or removed from *H* altogether. The computation of reduced transfer functions provides a convenient way to summarise network state from a subset of modes. Selection of modes by peak frequency can be useful as an alternative to integrating across the spectrum within specified frequency bands. In cases where a spectral peak lies close to the edge of a specified band, mode selection will allow the full contribution of that mode to enter the average without cropping its width to fit the band. The mode selection scheme can be tuned to fit the priorities of the research question at hand.

#### 2.1.4 Spatio-Spectral Eigenmodes

We define a Spatio-Spectral Eigenmode by the resonant frequency, damping time and transfer function of a single component in the modal decomposition of an MVAR model.

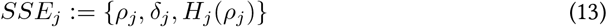

derived from a given eigenmode *j* in the eigendecomposition above. The transfer function of an SSE is evaluated only at the peak frequency of the mode in question. We will often split the total set of SSEs to explore the properties of a subset defined by permutation, frequency range or both.

### 2.2 Validation in simulated data

To illustrate the MVAR modal decomposition and Spatio-Spectral Eigenmodes we explored a single simulated dataset and a group simulation designed to exhibit realistic inter-run or inter-participant variability. The simulation scheme is summarised in figure 1A and described in detail in section 4.4. Briefly, the simulated activity in this network contained two resonances with pre-specified spatial and spectral structures. Two modes with distinct spectral structures were defined by direct pole placement and used to generate time-courses which were projected into a 10 node network structure. 20 realisations of 300 seconds in duration were simulated from this network structure.

**Fig 1.**
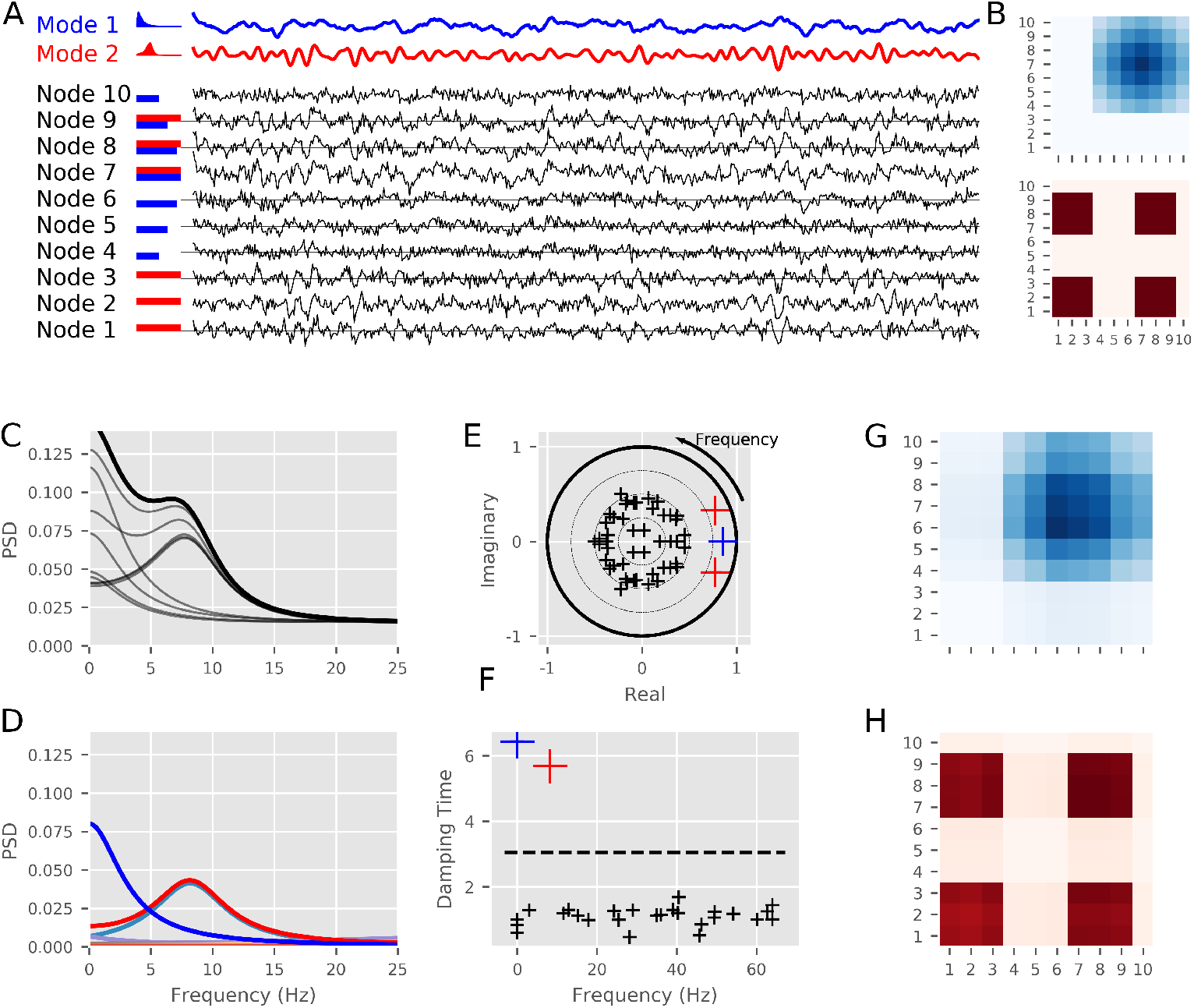
The simulations and modal decomposition for a single realisation of the simulated data. **A:** Summary of the simulation. ten nodes are generated from linear combinations of two modes with different spectra. The modes are shown in blue and red with the nodes in black, horizontal bars indicate the weighting of the two modes into each of the 10 nodes. **B:** Summary of the true network structure of the two modes. **C:** The power spectral density from ten nodes. These capture the gradual drop in frequency from 0Hz and a peak around 9Hz split across the different nodes. **D:** The modal power spectrum for the node highlighted in black in C. The gradual slope from 0Hz and the 9Hz peak are clearly isolated as distinct resonances. The remaining modes have very low amplitudes with no clear peaks. **E:** *z*-plane representation of the modal eigenvalues (shown as crosses). The frequency of each mode relates to its angle as it increases counter-clockwise from (1,0) to (−1,0). Negative frequencies correspond to angles increasing clockwise from (1,0) to (−1,0). The two high-power modes from D are clearly visible as the modes with the largest magnitude (red and blue). The remaining modes have small magnitudes and have evenly distributed angles (black crosses). **F:** Damping-time of each mode as a function of frequency. The red and blue modes have significantly longer damping times than the null distribution (99% threshold shown as the dotted line). These relate to 0Hz and 9Hz resonances in the data. The remaining modes have short damping times indicating that the influence of these modes is very short-lived. **G:** Modal PSD matrix computed from the eigenvectors associated with the blue mode (via the transfer function). **H:** Modal PSD matrix computed from the eigenvectors associated with the red mode (via the transfer function).

#### 2.2.1 Modal decomposition of a single session

An example segment of simulated data with its generating modes is shown in figure 1A. A non-oscillatory (blue) and an oscillatory (red) source time-course is created and projected across a network to create 10 node time courses (black). The red and blue horizontal bars indicate the weighting of each model time-course into each of the 10 nodes. The “true” network matrix containing the structure of each mode is shown in figure 1B. The time-series were described with an order-5 MVAR model fitted across the whole 300 second simulation. The Fourier based transfer function and power spectra were computed from this model and the spectrum of each node is shown in figure 1C. These spectra show the contributions from the two modes across the ten nodes. Some nodes contain signal from mode one (e.g. node 1), mode two (e.g. node 4) or both modes (e.g. node 7). Next, the modal decomposition was computed and the PSD con for node 7 (shown in black in figure 1C) is shown in figure 1D. Whilst node 7 contains contributions from both modes which are mixed in the Fourier-based analysis (figure 1C), these are clearly split into separate peaks (blue and red) in the modal power-spectrum (figure 1D).

The frequency *ρ* of each mode was computed directly from the eigenvalue *λ* of the fitted MVAR model. A *z*-plane plot of the eigenvalues of the decomposition (figure 1E) reveals that the 1*/f*-type mode is represented by a single real-valued mode (blue cross), in contrast the 9Hz mode is modelled by a complex conjugate pair of modes (red crosses). In the z-plane, frequency is represented by the angle of the complex eigenvalue, whilst the magnitude of the mode is its distance from the origin. As a more intuitive alternative, we show a scatter plot with individual modes with peak frequency on the x-axis and damping time *δ* on the y-axis (figure 1F). The damping time indicates how quickly an oscillation in that mode would be extinguished, longer damping times indicate that energy in the oscillation will dissipate more slowly. The damping time plots emphasise the dynamically important modes with long damping times whilst the frequency can be directly read out from the x-axis. In addition, the calculated damping time threshold for the simulation run is shown as a dotted line, demonstrating that the two relevant modes are easily separable from the background (Supplementary figure S.1 contains this data for all 20 realisations). Finally, the network structure of each of the two modes can be reconstructed from their modal transfer functions, computed from the relevant eigenvectors. Figures 1G and H show the Modal PSD of modes 1 and 2 respectively, each evaluated at its peak frequency (as determined from the respective eigenvalue). These reproduce the ground-truth structure shown in figure 1B.

#### 2.2.2 Modal decomposition of group-level networks

Next we examined how the Fourier band-integration and SSE approaches can describe oscillations with between subject variability in peak frequency. We computed 20 realisations (representative of 20 participants) of the simulation in figure 2 with varying peak frequency and amplitude in peak 2 whilst keeping the network structure itself static. Figure 2A shows the spectra of node 7 across the realisations of the simulation. The alpha peak frequency has a uniform +/−2Hz variability across realisations (gray lines represent individual subject); the group average can be seen as the solid black line. As in the single case, node 7 contains a contribution from both oscillatory networks; showing a 1*/f* type slope and a peak at around 9Hz. The simulated variance in peak frequency can also be seen clearly in the Fourier spectra shown in figure 2B; the frequencies-of-interest are highlighted in red and blue. The Fourier spectra captures the average features well, but the use of pre-determined frequency bands leads to clipping at the edges of some peaks. In addition, we can see contributions from the 0Hz peak influencing the shape and magnitude of the PSD around the 10Hz oscillation. Whilst adapting the frequency band of interest to the individual peak frequency could reduce the effect of peak clipping in the Fourier integration approach, it is harder to reduce interference between oscillations with overlapping spectra. As an alternative, the modal PSDs are shown in figure 2C. These are computed from the reduced transfer functions using poles selected by their damping time and driving frequency. In contrast to Fourier integration, this approach extracts single-peaks which vary depending on specific frequency content of the data.

**Fig 2.**
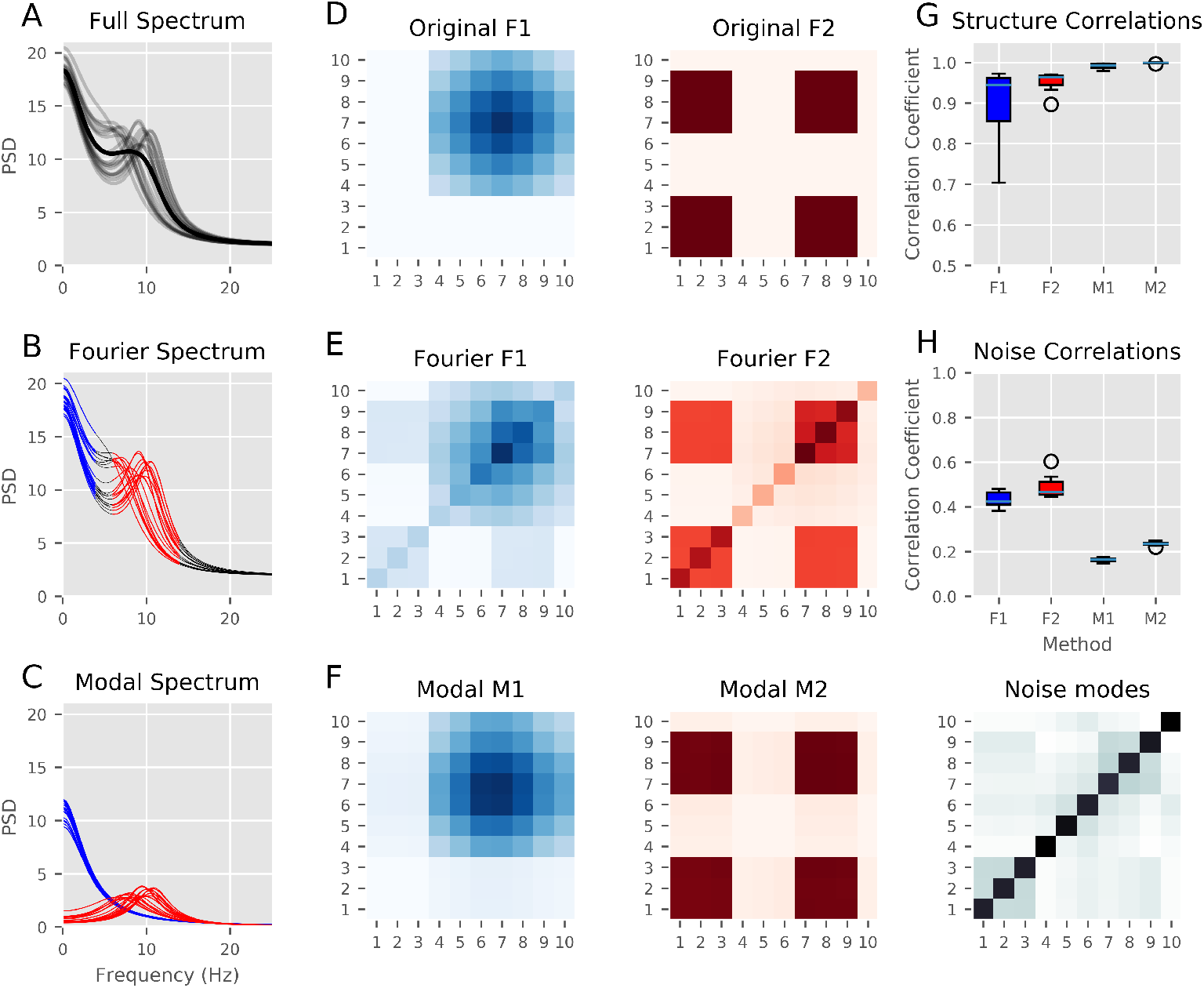
Power spectra and networks from the group simulation. **A:** The PSD of node 7 for all 20 realisations with variable spectra (gray) and the group average (black). **B:** Fourier-based PSD of node 7. The PSDs are split into pre-specified ‘low’ and ‘high’ frequency bands (in blue and red respectively). Though these capture the features around each frequency, they do not account for either individual variance in peak frequency or overlap between adjacent frequencies. **C:** Modal-based PSD of node 7. The modal spectra identified by thresholding the damping times of each mode of the modal decomposition and assigning each mode to its closest band (low in blue or high in red). The modal spectrum contains a single peak per mode and allows for variability in peak frequency between participants. **D:** Original network structure matrices. This figure shows the ground truth for the simulations generated in figure 1. **E:** Fourier-based network structure reconstruction. The network structure estimated from the Fourier-integration approach based on the bands in 2B, this captures the main structure with some interference from the adjacent frequency band. **F:** Modal-based network structure reconstruction. The network structure estimated from the modal bands seen in 2C. Here, the two spectrally distinct networks are properly resolved and there is little interference between the two. The diagonal structure in 2E is contained within the noise modes that did not survive the thresholding. **G:** The level of correlation (across the twenty realisations) between the ground-truth network structure and the network structure extracted for each individual run and both the Fourier and Modal analyses. The modal matrices show a much larger correlation with the ground truth than the Fourier-integration derived matrices. **H:** The correlation between the noise modes and the ground truth and Fourier network structures. The Fourier-integration matrices have a large correlation with the diagonal structure which is not explicitly associated with either of the simulated structures.

The group simulation used the network structure defined in figure 2D, with the network connectivity pattern driven by the 1*/f*-type signal on the left and the simulated alpha oscillation on the right. The network structure estimated by the Fourier frequency band integration approach captures the core features of the ground-truth simulations, but show spectral ‘leakage’ between the two underlying network patterns (figure 2E). The high and low resonances overlap in the frequency axis leading to low frequency content leaking into the high frequency integration window and vice-versa. In contrast, this mixing is absent in the modal estimation (figure 2F) which is able to separate the contribution from each pole to all frequencies and tune itself to variance in individual peak frequency. Both methods achieve a high (r *>*.9) correlation between true and estimated network structure across with the whole network and all realisations, however the modal networks are much more tightly clustered close to 1 (figure 2G). In addition, the noise network estimated from the residual modes correlates between r=.4 and r=.5 in the case of the Fourier-integration estimated networks whilst the same correlation in the modal networks is much lower (around .2) (figure 2H).

### 2.3 Oscillatory networks in MEG data

We next explored the frequency structure of oscillatory networks in the Human Connectome Project MEG data. We demonstrate that an autoregressive model can capture the frequency specific content of a whole-brain functional network using standard Fourier band-integration before moving to explore the SSEs. A summary of the whole analysis pipeline for the HCP is given in figure 3. Pre-processed MEG data were source localised to a 5mm grid throughout the brain using an LCMV beamformer before groups of voxels were combined into parcels based on the 78 cortical regions in the AAL atlas. The parcel time-courses were then orthogonalised to reduce leakage (details on the MEG processing are included in section 4.5). MVAR models were fitted to each recording session before their Power and Cross Spectral Densities were estimated using the Fourier-integration approach (Model fitting and validation is described in detail in section 4.6 and 4.7).

**Fig 3.**
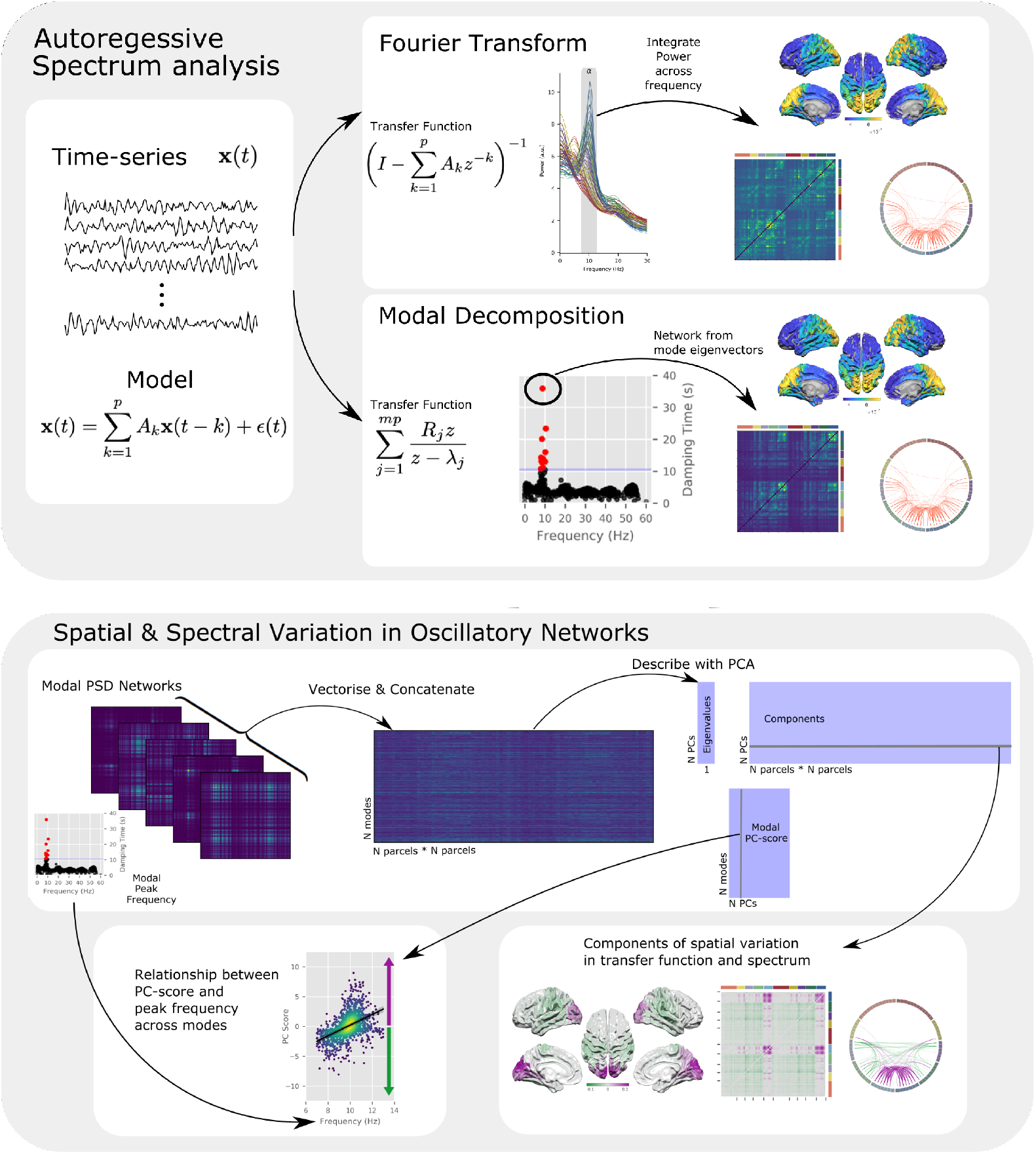
A graphical outline of the analysis procedures used in this paper. **Upper:** Summary of the procedures used to calculate the MVAR model and analyse the results using a Fourier (upper section) or Modal (lower section) approach. The Fourier-integration approach is used in figure 4 and the Modal decomposition is explored in example participants in 5 and at the group level in 6 **Lower:** Outline of the procedures used to take the modal decomposition of the MVAR model and compute spatial principal components each of which can explain variability in different frequencies within and between-participants. The group results of the PCA analysis are presented in figure 7 and a summary of the results of individual participants in figure 8.

**Fig 4.**
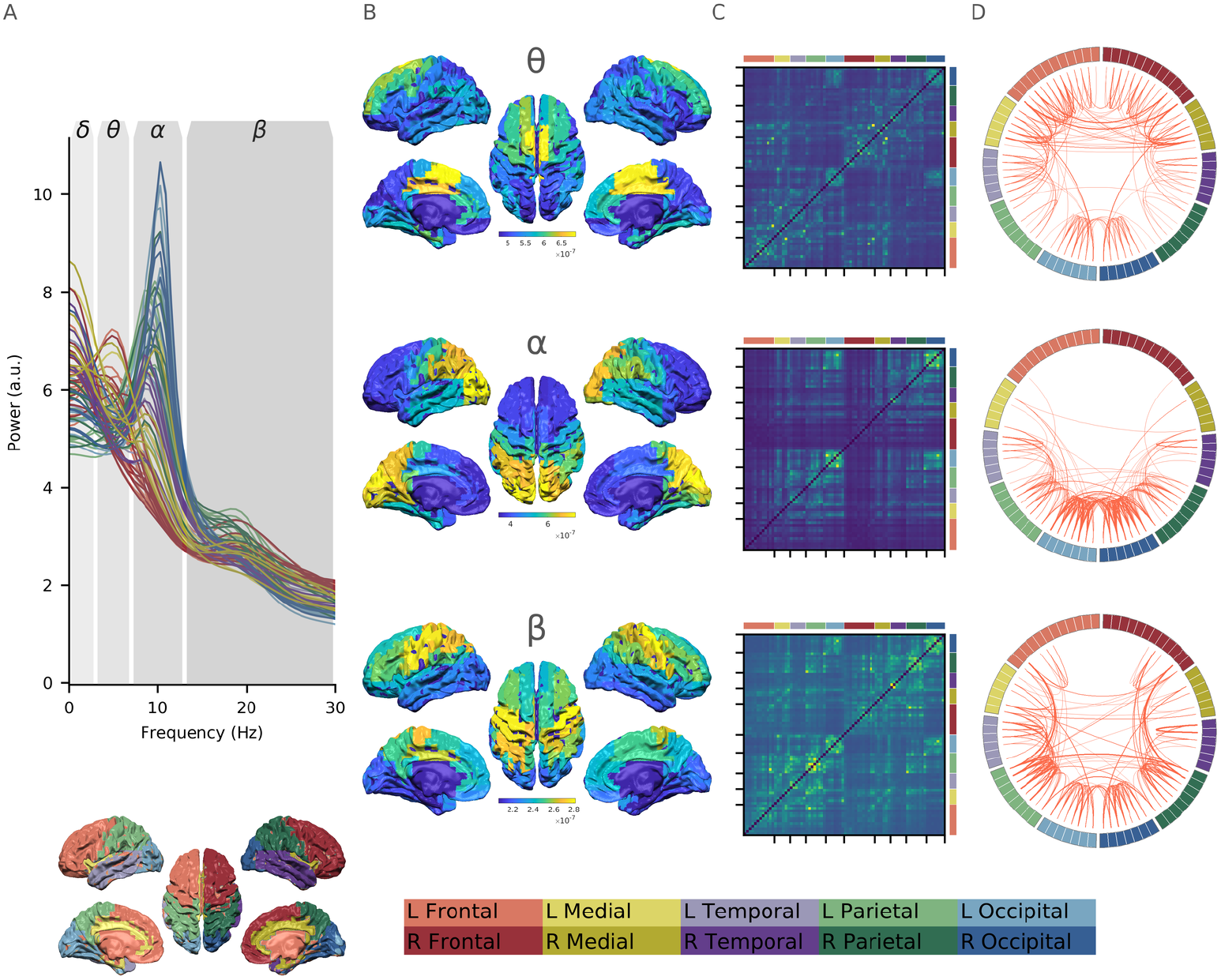
Fourier-based frequency-specific networks extracted from fitted MVAR parameters. **A:** The Fourier Power Spectral Density averaged across participants for each node in the AAL parcellation. The model captures a clear 1/f type trend across the spectrum as well as a distinct alpha peak. The diagram below shows the colour code for each region of the AAL atlas. **B:** Surface plots showing the average PSD for in each cortical parcel within the theta, alpha and beta bands. **C:** Network matrices showing the average CSD between parcels within the theta, alpha and beta bands. **D:** Circular network plot showing the average CSD between parcels within the theta, alpha and beta bands. Region colouring is shown in the legend at the bottom of the figure: red: frontal, yellow: medial, purple: temporal, green: parietal, blue: occipital. Lighter colours refer to the left hemisphere (and are on the left of the network matrices and circular plots) and darker colours refer to the right hemisphere (and are on the right of the network matrices and circular plots).

The topography of MEG functional networks vary as a function of frequency [32–36]. As a result the spectrum is typically split into a set of independently analysed frequency bands. Our autoregressive model fits were able to capture frequency specific power distributions and network structure within these *a priori* defined bands. The average Fourier derived PSD (across participants) from each node of the AAL can be seen in figure 4A. Overall, each node shows a 1*/f* trend with the strongest oscillations visible in the alpha band. Frequency-specific source topographies are shown in the remaining columns of figure 4. These maps were computed by averaging the MVAR PSD estimates across participants within *a priori* frequency bands. Each panel includes source-space images containing the diagonal of the PSD matrix within the frequency band as well as a network matrix showing the off-diagonal Cross-Spectral Densities (CSD) and a circular connectivity plot showing the network connectivity based on the CSD. The circular connectivity plots show the connections whose magnitude falls within the top 5% of the off-diagonal CSD distribution for each frequency band. Details of the labelling and colouring of each cortical region can be found in the figure caption.

The 3–7Hz theta band has strongest power in medial prefrontal regions with connectivity including connections with the parietal and occipital cortex. In contrast, alpha (7–13Hz) power is strongly localised to occipital cortex with strong connections within the occipital region and between the occipital and temporal regions, with a smaller number of parietal connections. Beta power (13–30Hz) is predominantly seen in the bilateral motor cortices with a broad range of connections. Overall, these results suggest that MVAR model based Fourier frequency-domain power and connectivity estimates are able to represent whole-brain functional connectivity patterns in line with expectations from the literature [37].

### 2.4 Spatio-Spectral Eigenmodes capture individual variability in oscillatory networks

The Spatio-Spectral Eigenmodes (SSEs) provide an alternate description of network power spectra based on their characteristic frequency and network structure. The modal decomposition was computed for the MVAR model of each recording session yielding *mp* SSEs (in this case an MVAR model over 78 regions with model order 12 yields 938 SSEs), though only a minority of these reflect dynamically important structure in the data. Non-parametric permutations were used to split the full set of SSEs into an included set of dynamically relevant functional modes and an excluded set of modes whose damping times are not distinguishable from chance in this dataset (see section 4.9 for details). Figure 5 shows the Fourier power spectrum and SSEs for four individual participants (in rows). The first participant has a strong alpha peak at around 11Hz (Fourier power spectrum shown in column A) which is well represented by the SSEs with long damping times (scatter plot of SSE damping times by peak frequency in column B). The SSEs in red are included in further analyses having survived the non-parametric permutations. The power spectrum from full and included set of SSE (black and red lines in column C respectively) again indicate that the included SSEs do capture the prominent oscillations in the full power spectrum. Furthermore, the included SSEs have a bilateral spatial distribution and network structure around the occipital pole with a bias toward the the right hemisphere (included SSE network connectivity matrix and surface plot in columns D and E). The second participant has a similar alpha peak in the power spectrum with a slightly lower peak frequency. In contrast to the first participant, this participant’s alpha is broadly distributed around bilateral occipital and parietal regions. The third participant has a small alpha peak corresponding to significant SSEs with relatively short damping times compared to participants 1 and 2. A single SSE in the beta band survives the permutation scheme and contributes to a diffuse power and network structure between occipital, parietal and motor cortex. Finally, example participant 4 shows two separate alpha peaks in two different brain regions as shown by the 8 and 10Hz peaks in the power spectrum and SSE damping time plots. The average spectrum shows a prominent, relatively low frequency alpha peak which is will described by the significant SSEs. Similar to participants 2 and 3, this participant has a relatively diffuse alpha power distribution across occipital and parietal cortex.

**Fig 5.**
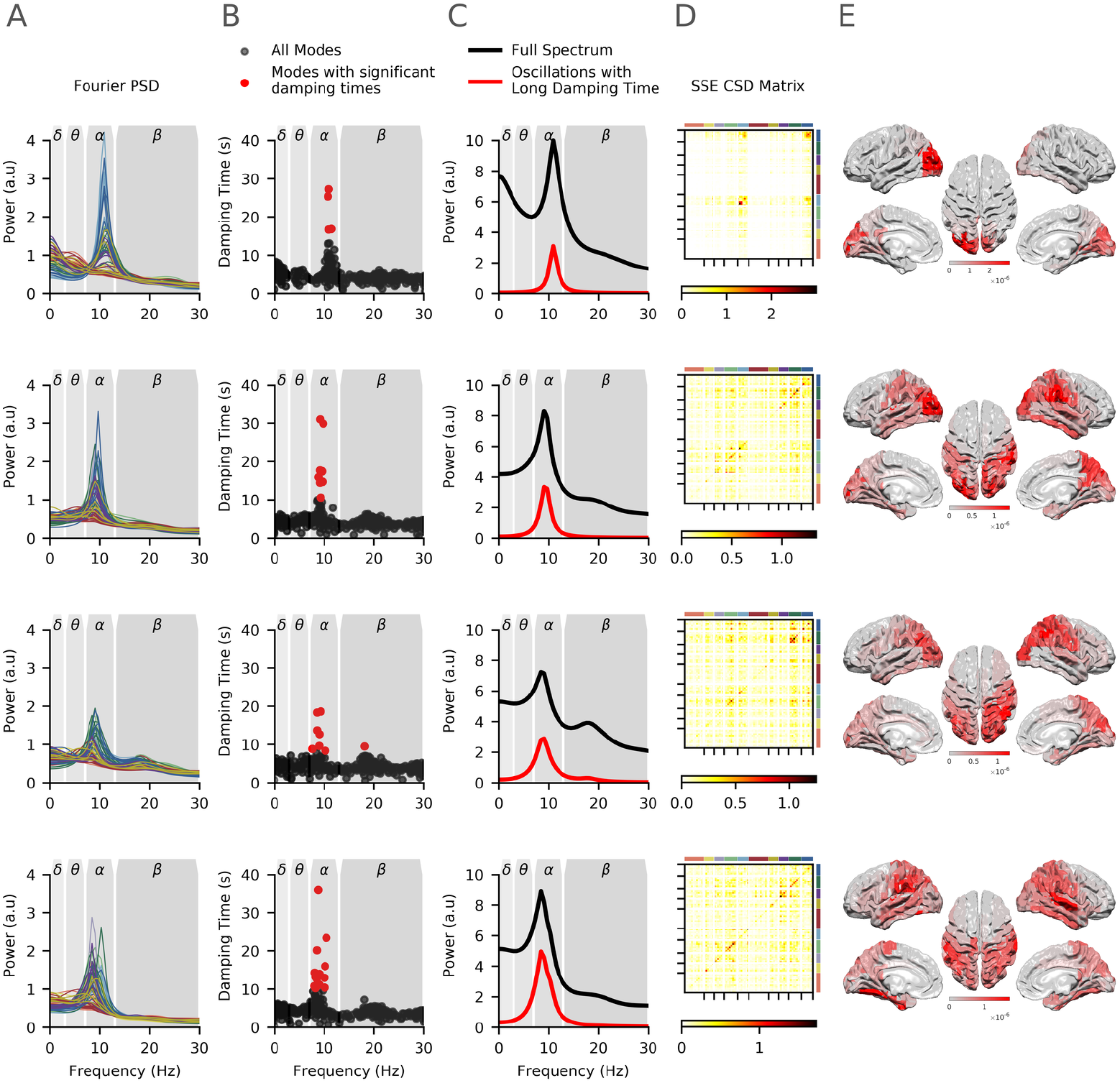
Spatial and spectral variability illustrated by data from four example participants. Each row contains the Fourier power spectrum and modal decomposition for that participant. **A:** The standard Fourier power spectrum, coloured lines indicate brain regions following the colour code in figure 4A. **B:** The damping times of the modal decomposition as a function of mode peak frequency. All modes are shown with dynamically important modes shown in red. **C:** The power spectrum averaged across all brain regions computed using the Fourier method (black) and the reduced spectrum computed from the modes surviving the permutation tests is shown in red. **D:** Network connectivity matrices computed from the reduced modal transfer function (corresponding to the red line in C). **E:** Spatial distribution of power from the reduced modal transfer function corresponding to the diagonals in D.

### 2.5 Group variability in alpha peak frequency

To describe the oscillatory frequency content across the whole brain and group, the damping times of all modes across the full HCP dataset are plotted as a function of peak alpha frequency in figure 6A. The included sets of SSEs (as identified by non-parametric permutations) are indicated in red with the remaining SSEs in black. As in the individual cases, the modes with the longest damping times occur around the strongest peaks in the Fourier spectrum (figure 4A). The majority of these (for the present eyes-open resting state data) lie within the traditional alpha range with a smaller number in the delta, theta and beta ranges and a single mode above 30Hz. In contrast, the excluded SSEs are relatively distributed across the whole frequency range. Figure 6B shows the average power spectrum across all regions and participants (black) and the spectrum reconstructed from only the significant SSEs (red). The relatively small number of SSEs preserve the prominent oscillations in the signal at the group level suggesting that the permutation scheme is successful in extracting the SSEs related to the largest resonances in the system. The predominance of alpha in the surviving SSEs reflects the prominence of the alpha rhythm in power spectra across eyes-open resting state MEG scans. This is a static power spectrum estimate across the whole duration of each scan. As such it is possible that individual variability in these alpha peaks are driven by temporal dynamics as well as oscillatory amplitude, similarly there may be transient bursts in other frequencies that are not detected in the average spectrum [38].

**Fig 6.**
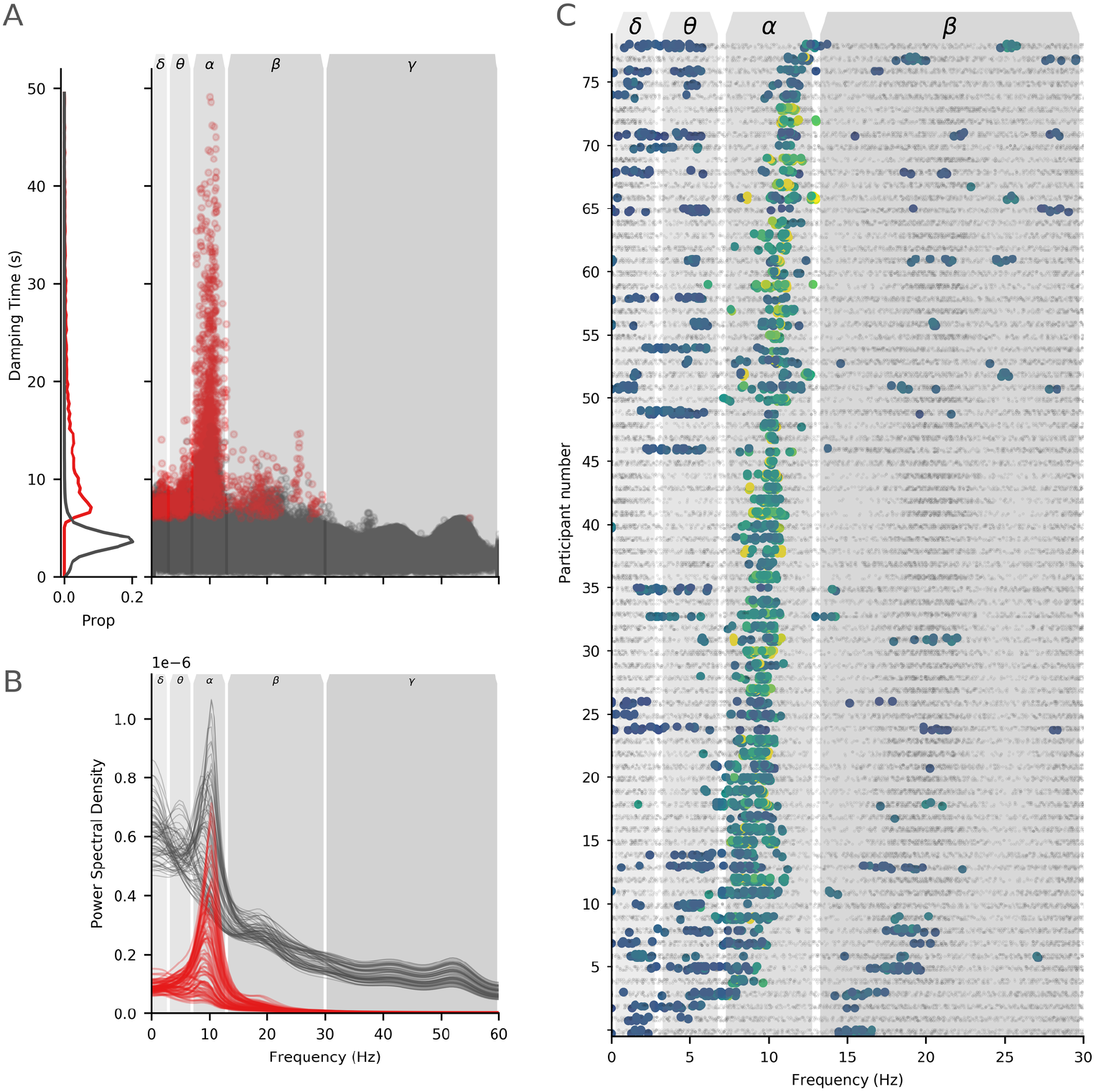
Assessment of modal poles in individual participants. **A:** Plots of damping time against frequency for all poles, for all participants. Poles coloured in red survived the non-parametric individual subject thresholding and were carried through to later analyses. The left hand plot shows the normalised distributions of poles which survive (red) and do not survive (black) the thresholding procedure against damping time. **B:** The power spectrum of all modes (black) and modes surviving the non-parametric permutation scheme (red) across all datasets. Each region in the AAL atlas is shown as an individual line. **C:** Plot of the modes for all participants sorted by peak alpha frequency. Each row on the y-axis shows an individual participant and each dot in the graph an individual mode (all three runs for each individual are combined in one graph). Below threshold modes are shown as small black dots whilst modes which survived thresholding are shown as larger dots in colours. The x-axis shows the frequency of the mode and a set of canonical frequency bands are shown in gray boxes.

Across all participants the frequency distribution of SSEs provides a straightforward summary of the spectral variability in the HCP dataset. (figure 6C; surviving modes are shown as blue-green dots in rows) though the three runs within each participant are quite consistent. The rows of figure 6C are sorted by Individual Alpha Frequency (IAF; derived from an average frequency of SSEs between 6 and 14Hz, weighted by damping time) showing the variability across participants. The majority of surviving modes fall within the traditional 7–13Hz alpha band. 76/79 participants have at least one significant mode within alpha although peak frequencies are widely variable across participant (median peak frequency of alpha modes across participants range from 7.4Hz to 12.9Hz; mean equivalent from 7.4Hz to 12.9Hz). The participants with the lowest and highest IAFs lie very close to the boundaries of the standard 7–13Hz frequency range. Though the peak frequency lies within these bounds and are therefore well represented by the SSEs, the standard alpha band does not contain the full width of the oscillatory peak for these participants. As such, an analysis that imposed a strict cut-off would likely clip the edges of these alpha peaks leading to potential distortions or misrepresentation of the spectral content. A relatively small number of oscillatory modes in some participants occur within the delta, theta and beta bands.

### 2.6 Alpha peak frequency varies between occipital and parietal cortex

Each SSE is a property of the whole brain rather than a single region or ROI, allowing it to represent the distribution of an oscillation across space and network connections. The spatial variability in the SSEs PSD networks whose peak lies within the 7–13Hz alpha range was described by a small number of components in a Principal Components Analysis (PCA). The components of PCA analysis describe the axes of spatial and network variability across modes whilst the Principal Component (PC) scores indicate the extent to which a particular PC component is expressed within a given SSE. The reproducibility of each PC was evaluated by split-half correlations, this indicated that the first two components were highly replicable across halves of the data (the validations for the PCA analysis are described in detail in section 4.10 and the results are shown in the Supplementary Material in section 9.4)

Crucially, this PCA was computed on the spatial network structure of alpha SSE without knowledge of the corresponding resonant frequencies. We next quantified the correspondence between the spatial content and the peak frequencies of the SSEs across participants. A Bayesian regression was used to assess the extent to which peak frequency can be used to predict PC score across SSEs. There is large between subjects variability in alpha peak frequency (figure 6C) so we include each participant as a random effect term in the model. This allows us to directly quantify the between subject variability in peak frequency and explore whether there is a consistent relationship between network structure and peak frequency across the dataset even where the absolute peak frequency itself is variable. Further, to assess whether a term in the model provides good out of sample predictions, we used a Leave-One-Out (LOO) cross-validation procedure. Details of the Bayesian model inference and validation are described in section 4.11.

This PCA analysis was repeated for SSEs lying within the theta and beta bands. SSEs within these bands reflected the expected spatial structure of theta and beta activity (details in section 9.5). As relatively few theta and beta SSEs survived thresholding by non-parametric permutation, we did not go on to perform a detailed investigation of the their spatio-spectral covariation. The small number of theta and beta modes surviving permutation reflects the predominance of alpha oscillations in resting state recordings. The distribution of modes reflects the content of the signal, so we would expect to find more theta and beta modes in the task evoked data.

The first principal component in the alpha band (PC1: 23.0% variance explained, *r* = .94 average split-half correlation) relates to the average power across an occipital network similar to the standard alpha network. The component values for PC1 have the same sign in all brain regions indicating that changing PC-scores will act to increase or decrease power across this whole network. In other words, SSEs with a positive score in PC1 will have a high power across this distribution, whilst SSEs with a negative score will have low power across the whole brain (figure 7 PC1). For the first principal component, the difference in LOO scores between the intercept-only and full models was −9.7 (SE: 5.3), indicating that there is insufficient evidence to conclude that PC-score (corresponding to overall power) is related to peak frequency in this component.

**Fig 7.**
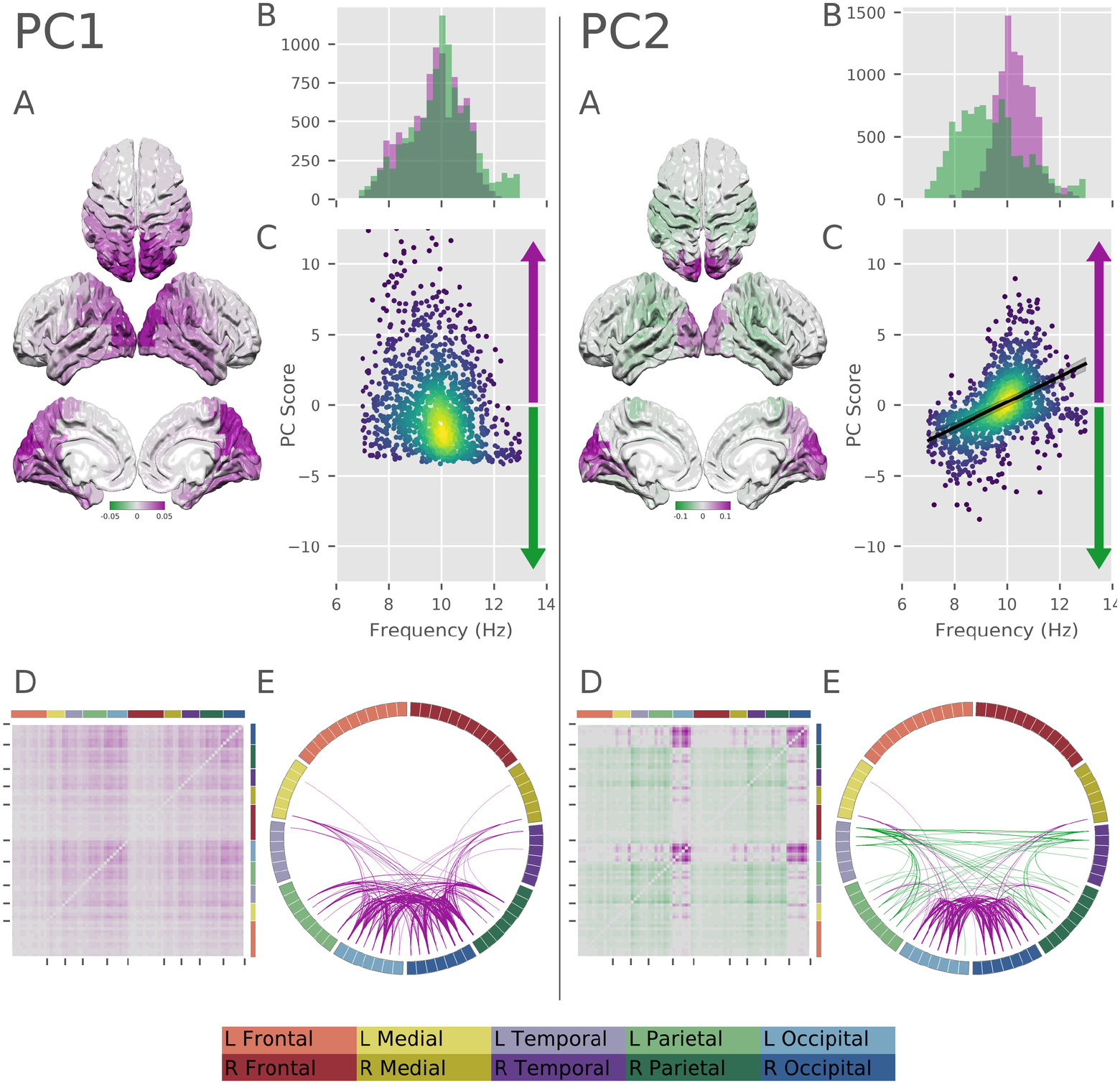
Results of the PCA decomposition of the modal analysis for PCs 1 and 2. For each component we show: **A:** a surface plot of the component structure across nodes in the AAL atlas. **B:** histograms showing the distribution of SSE frequencies for SSEs with positive (purple) and negative (green) scores. **C:** scatter plot showing the relationship between SSE peak frequency and PC-score. For PC2, this also contains the regression line quantifying the modelled relationship. The frequency component of the model was not significant in PC1. **D:** network matrix showing the component structure across network connections. **E:** circle plot showing the component structure across network connections (top 15% of connections shown).

The second alpha component (PC2: 12.7% variance explained, *r* = .43 average split-half reproducibility) contains a spatial gradient with the occipital pole at one end and parietal lobes at the other. Power at the two ends of this gradient are in counterpoint, a positive score for PC2 indicates high power in occipital lobes with suppressed power in parietal lobes and vice versa for negative scores. The Bayesian model was used to assess whether SSE peak frequency can be used to predict PC-score for PC2. In this case, the difference in LOO scores between the intercept-only and full modes was −102.6 (SE: 13.0) indicating that there is sufficient evidence to warrant assessing the full model. The frequency parameter in the full model had a central parameter estimate of 0.46, with a 95% CI of 0.40–0.52. This indicates that an increase of peak frequency of 1Hz would correspond to an increase in PC-score of around 0.46 of a standard deviation of the distribution of scores. In other words, increasing peak frequency across SSEs corresponds toward increased power in occipital cortex and decreased power in parietal cortex. The overall distribution of SSEs between the two principal components is shown in figure S.8.

#### 2.6.1 Within and between subject variability in alpha frequency

The distribution of SSE damping times as a function of frequency (figure 5) and the random effects term in the Bayesian linear model indicate that there is very wide individual variability in alpha peak frequency. Next, we visualise how this between subject variability interacts with occipto-parietal gradient in PC2. Figure 8 shows the distribution of SSEs with positive and negative PCs scores as a function of frequency for PCs 1 and 2. PC1 shows a wide distribution of peak frequencies between 7 and 13Hz (figure 8A). The distribution of frequency differences between SSEs with positive and negative scores are nearly completely overlapping (figure 8B).

**Fig 8.**
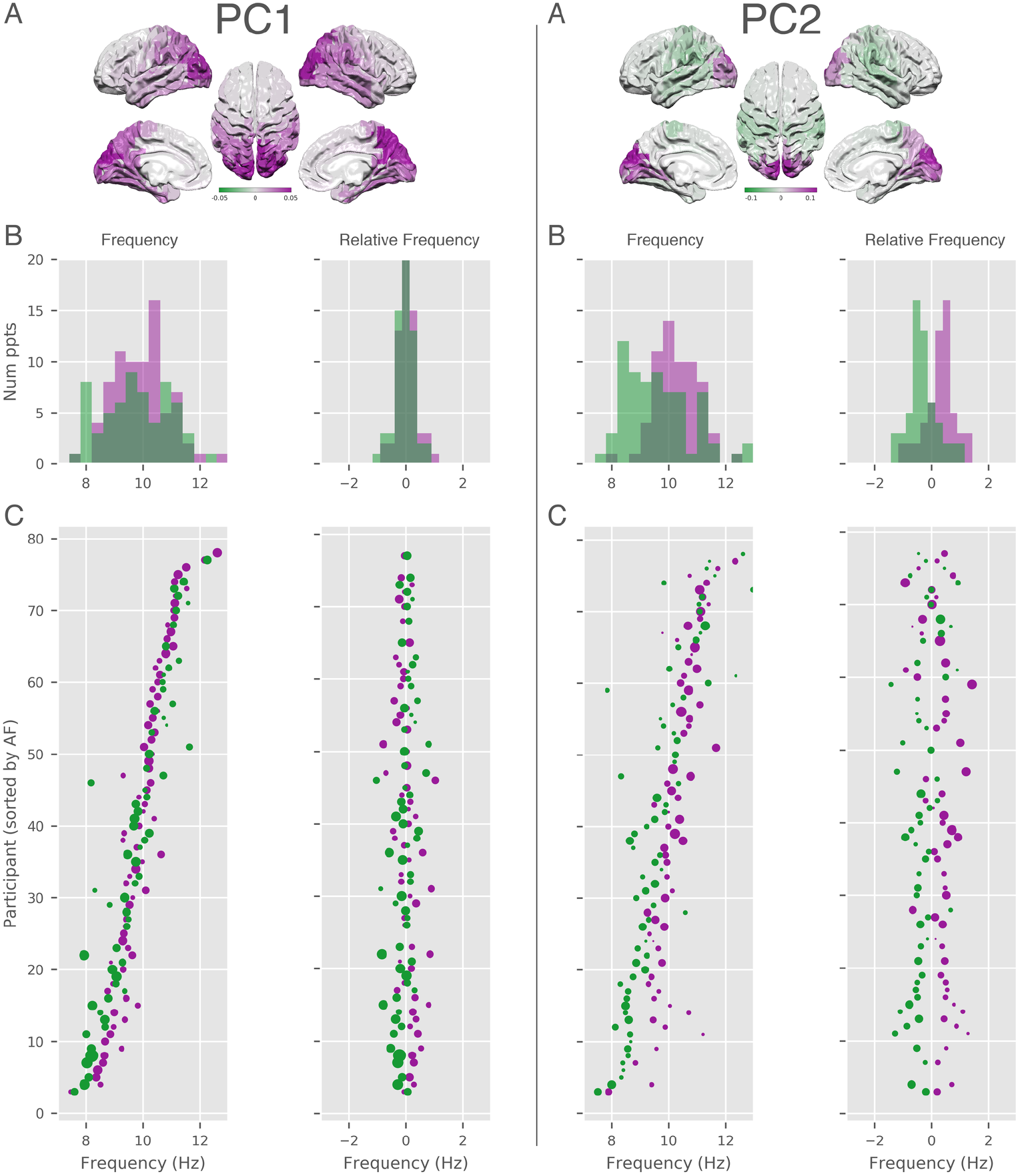
Correspondence between the PC scores and peak frequency of the surviving modes for PC1 and PC2. For each component we show. **A:** a surface plot of the component structure across nodes in the AAL atlas. **B:** Histograms of the SSE peak frequency for each mode split by positive (purple) and negative (green) score for the component. The left hand histogram shows the absolute mean peak frequency for each individual for the positive and negative scores. The right hand histogram shows the frequency of the components relative to the participant mean. **C:** Per-participant scatter plot of the SSE peak frequency for each mode split by positive (purple) and negative (green) score for the component. The left hand scatter plot shows the absolute mean peak frequency for each individual for the positive and negative scores. The right hand scatter plot shows the frequency of the components relative to the participant mean. The order of participants is sorted in the same manner as in figure 6 C.

In contrast, the distribution of PC2 frequencies shows a mean shift between positive and negative PC scores in both the absolute and relative distributions. Parietal alpha has, on average, a lower peak frequency distribution than occipital alpha. The two between subject distributions begin to overlap around 9Hz. Using the SSE methods to un-mix spatial and spectral variability we can see that parietal alpha SSEs occur between 7–11Hz across participants. The higher end of this distribution would otherwise be masked by the stronger occipital power at frequencies above 9Hz. A key source of this mixing is that variability in overall alpha peak frequency between subject is larger than the frequency difference between parietal and occipital alpha. Specifically, the overall alpha peak distribution ranges between 7 and 13Hz (range of 6Hz), though the relative difference between the two ends of occipito-parietal gradient is around 1Hz.

## 3 Discussion

Spatio-Spectral Eigenmodes defined from the properties of an autoregressive model provide a flexible representation of oscillatory brain networks with minimal prespecification of regions of frequencies of interest. We introduce the theory behind this approach and demonstrate its application in simulations and resting state MEG data from the Human Connectome Project. Firstly, we established that an autoregressive model is able to describe frequency specific functional networks in whole brain MEG data. The modal decomposition is then computed on these models to identify Spatio-Spectral Eigenmodes (SSEs). The resonant frequencies and damping times of these SSEs are shown to provide a simple summary of the oscillatory content of whole functional network. The spatial distribution and network structure of each SSE can then be explored through its contribution to the system transfer function and subsequently, power and cross spectral density. We utilised these properties to explore spatial and spectral variability in alpha oscillations. The SSEs expressed the between subject variation in individual alpha frequency and showed that, on average, alpha oscillations in the parietal lobe are lower frequency than those in the occipital lobe. Though this is a robust group effect, wide between subject variability means that the definition of ‘low’ or ‘high’ frequency are overlapping across individuals. A 10Hz oscillation could load onto the parietal lobe in one participant and the occipital lobe in another. The SSE approach separates these sources of variability by computing network structure and peak frequency simultaneously in each dataset. The SSE parameters then provide a convenient and intuitive representation of the spectral shape, spatial topography and network connectivity of neuronal oscillations.

### 3.1 Spatio-Spectral organisation of alpha networks

The oscillatory signatures of brain function which are measured during eyes open resting state are dominated by the alpha rhythm. The first spatial component of alpha power identified by our analysis describes variations in mean power in a network centred, on average, around medial-occipital cortex [16, 17]. The individual resonant frequencies of these SSEs support a wide previous literature showing that IAF estimates between subjects vary widely within and around the traditional 7–13Hz range [7, 8]. Crucially, this variability is functionally relevant and has been linked with a wide range of cognitive and clinical markers [39]. This presents a practical problem in that the estimation of spatial maps or networks in participants whose alpha peak lies close to these bounds. If the whole width of the alpha peak is not within the specified range parts of it will be cut off, leading to possible distortions in the estimated maps and networks. One solution to this is to tune the centre frequency or width of the frequency bands to the peak of each individual subject [7, 8], however this depends on the accurate quantification of the peak. Here, we show that the parameters of MVAR-derived SSEs can overcome some of these limitations by characterising individual spectral peaks without pre-filtering data into frequency bands or locations of interest.

The second PC of spatial variability in alpha SSEs shows a distinction between networks loaded onto occipital or parietal cortex. On average, SSEs with high scores towards the occipital end of this component tend to have higher frequencies than those centered at the parietal end (10Hz compared to 8–9Hz). This split between low and high frequency alpha is similar to the low and high bands (typically defined around 8.5 and 10Hz in the literature) are suggested to reflect separate cognitive functionality [7]. Though this difference is strong on the group level, the frequency distributions of occipital and parietal SSEs are overlapping, suggesting that an oscillation of 10Hz could correspond to one participant’s low frequency parietal alpha and another participants high frequency occipital alpha. A small minority of participants in this study show the reverse effect, with higher frequency oscillations in parietal rather than occipital cortex. The difference between occipital and parietal alpha when both are present within an individual (around 1Hz) is substantially smaller than the range of alpha peak frequencies across participants (7–13Hz). As such, the full range of frequency variability in these regions is only visible with an analysis approach that can simultaneously deconvolve spatial and spectral variability.

The paper presents an exploratory analysis of a publicly available, eyes-open resting state dataset with the aim of characterising the structure and variability in oscillatory networks. Whilst we can show that alpha oscillations have these spectral and spatial distributions across a large dataset (from the Human Connectome Project), without an experimental task or prior hypothesis we do not make strong claims about its functional interpretation based on these analyses. To guide future research, we propose three potential interpretations of the distinction between occipital high-alpha and parietal low-alpha found in PC2. Firstly, these rhythms may reflect spatially and functionally distinct generators of alpha [20]. Occipital alpha is thought to represent the locus of visual attention [40] whilst, parietal alpha has been linked with attentional processing and is suggested to exert top-down control of visual alpha depending on attentional state [20, 41]. Our results show this distinction between occipital and parietal alpha may be present in resting-state data and that these alpha sources are additionally separated by peak oscillatory frequency. A second possibility is that the second PC identified in our analysis represents a continuous gradient of oscillatory behaviour between occipital and parietal cortex. Similar gradients in structural and functional MRI data have been proposed as an organising principle of the brain [42], PC2 may then represent a occipito-parietal gradient organising alpha oscillations. A related idea is that PC2 could reflect an aspect of the posterior to anterior alpha travelling waves [43]. Finally, the parietal end of PC2 may represent the sensori-motor Mu rhythm rather than a distinct parietal alpha source. The Mu rhythm peaks over sensorimotor cortex and has a similar frequency but distinct waveform shape to occipital alpha [44] Future research in this area using task-related data could distinguish between these hypotheses.

### 3.2 MVAR models: Parameterisation & Limitations

The Spatio-Spectral Eigenmode decomposition method is dependant on a good estimation of the power spectra of the system via the underlying MVAR model. In turn, the estimation of the PSD is dependent on adequate selection of the hyper-parameter of the MVAR model: the model order (p) and the sample rate of the data [45]. In the current work we downsample the source time-courses to 100Hz and use a model order of 12. This provides a good trade off between the high spectral resolution arising from high model order and straightforward model estimation from low model order (see SI 9.3 for further details). Further, as autoregressive models will always fit the entire spectrum from zero to Nyquist, the low sample rate ensures that the spectrum fit focuses on the physiological range of interest. Though these parameters, give a good fit in this instance, it is not guaranteed that they will generalise to novel datasets and appropriate diagnostics must be performed in these cases.

The modal-form of the transfer function has a spatial constraint; a single SSE is associated with a rank-1 network structure. More complex network structure is described through a combination of SSEs. This is mathematically straightforward as the transfer function can be summed across modes, yet the method for identifying which modes to combine must be tuned to the application in hand. The linear summation of modes is only equal to the full Fourier model at the level of the transfer function. Though properties such as the PSD matrix can be defined from a single SSE, the summation of these modal-PSD matrices will not necessarily equal the Fourier equivalent. Here, we explore the spatial and spectral properties of PSD matrices across many SSEs without directly summing them. Other applications may wish to combine these SSEs at the level of the transfer function for each data recording prior to group analyses. Finally, the modal cross-spectral densities used in the network analyses represent both the shared power and phase-locking between each pair of nodes. A richer representation of connectivity could be gained by using coherence or directed transfer function based metrics rather than the CSD, however the normalisation of these measures is difficult with the rank-1 matrix structure limitation in SSE analysis. We are continuing work into these issues and the wider picture of how the SSE decomposition and modal transfer function relates to standard power spectrum and connectivity measures.

### 3.3 Relation to other decompositions

The decomposition of autoregressive models of univariate EEG time-series into their natural frequencies, damping times and transfer function contributions has a long history [46–48]. Recently, the computation of natural frequencies and damping times has been generalised to multivariate autoregressive models [23]. We link these multivariate parameters to the system transfer function via Gilbert’s Realisation [30, 31] leading to the definition of the Spatio-Spectral Eigenmodes.

There are several mathematically related approaches in the literature. In particular the method in this paper are closely related to techniques for modal analysis which have widespead use in engineering. Firstly, SSEs are closely related to the Principal Oscillatory Patterns and Principal Interaction Pattern analyses of autoregressive models originally developed for analyses of climate systems [22]. The peak frequencies and damping times from the eigenvalues of these analysis have previously been used to investigate EEG recorded during epileptic seizures [49]. Next, a Hankel matrix can be used to identify a state-space parameters and permits a modal decomposition to identify mode frequencies and damping times (for example the Eigensystem Realization Algorithm; [50]). Decompositions of the Hankel matrix have been previously applied to explore the frequency modes of epileptic seizures [51]. Finally, the Dynamic Mode Decomposition (DMD) represents oscillatory dynamics via Koopman modes [52]. It is optimised for image-type datasets where there are more regions or channels than time-points in a dataset and has previously been applied to fMRI [53] and ECoG [54, 55] recordings. The application of these methods and their deeper mathematical relationship is a point of active research in the Neuroscience and the wider dynamical systems literature.

A range of conceptually related methods look to isolate oscillatory activity in electrophysiology data using linear spatial filters (see [56] for a review). Unlike the approaches above, these typically involve computing a frequency or time-frequency spectrum across the dataset and before carrying out the decomposition, often using using PCA, ICA or related techniques. The spectrum estimation and decomposition stages may be carried out and optimised separately. These decompositions tend to be relatively unconstrained in the frequency domain, the resulting component power spectra can be comprised of complex, multi-modal shapes which may be challenging to interpret as clear oscillatory signals.

## Conclusion

We have shown that a modal decomposition of MVAR parameters can be used to simultaneously estimate spatial and frequency structure within human resting state MEG data. In the SSE framework, brain networks are decomposed into oscillatory signals on an individual whole-brain basis with minimal pre-specification and averaging. Using this method, we have demonstrated that multiple, spatially overlapping, sub-networks exist within the normal alpha band activity. Detailed within-subject networks can be identified despite large between-participant variance. These structure captured by the SSEs can be used enhance investigation into how individual oscillatory phenotypes relate to individual difference in cognitive and clinical states.

## 4 Methods

### 4.1 Derivation of the modal form of the transfer function

The transfer function describes the filtering carried out by a linear system as it transforms an input into an output. In this case our input is a noise process defined by the residual covariance of a fitted autoregressive model and the output is the frequency transformed data.

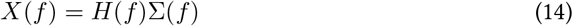

In the main text, we limit values of *z* to the unit circle (where |*z*| = 1) as these can be directly related to oscillatory frequencies *f* and the discrete time Fourier transform. In the following derivation we generalise this to evaluate *H* across any point in the *z*-plane by including a magnitude term *M* in the definition of *z*.

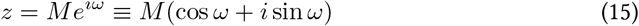

Where *ω* = 2*πf* Δ*t*. When *M* = 1, this is equivalent to evaluation in terms of oscillatory frequency *f*.

This transfer function can be written in several forms depending on the context. In autoregressive spectral estimation, this state-space form is commonly used (see equation 5 in the main text)

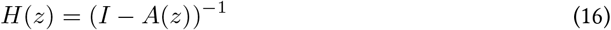

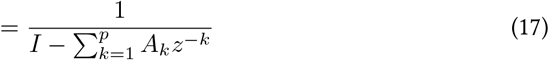

We compute the transfer function using *A* as we start with a simple autoregressive model with no moving average component, inputs or observation equations. The complete state-space form would include input, output and feed-forward components which we omit here for simplicity. For our autoregressive model, the numerator of the transfer function is 1 and the denominator is a polynomial series of the model parameters under a complex valued z-transformation. The denominator can be expanded to show the full polynomial form.

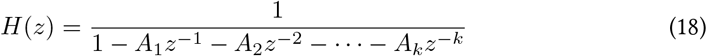

Here we see that the transfer function of a system is the fraction of two polynomials defining how the system transforms inputs into outputs. We can evaluate *H*(*z*) around the unit circle (*z* = *e*^−*i*^2^*πft*^) where values of *z* directly correspond to different frequencies. The output of *H*(*z*) can then be used to compute a power spectrum as seen in equations 2, 5 and 6. Though the transfer function can be evaluated for any value of *z*, the magnitude of *H*(*z*) around the unit circle, and therefore the estimated power spectrum, are dependant on the roots of the polynomial *A*. The roots of *A*(*z*) are called poles (*λ*) and they define coordinates in the *z* plane at which *A*(*z*) will evaluate to zero, driving the value of *H*(*z*) to infinity. We can rewrite equation 18 in terms of the poles *λ*.

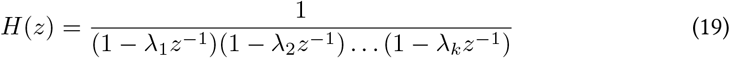

The value of *H*(*z*) decreases as the distance of *z* from each pole increases. Where there are multiple poles in a system, the value of *H*(*z*) at a given point is dependant on all of the poles, though as the influence of a pole falls off rapidly with increasing distance the closest poles to a point *z* have the greatest influence. As such, the value of the transfer function at a given point *z* is linearly dependant on the distance of *z* from this set of poles, providing an alternate formulation of *H*(*z*) (assuming no repeat poles).

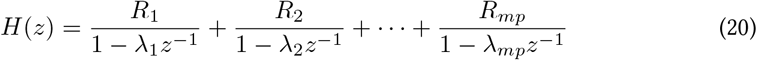

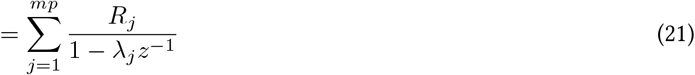

Here, *R_j_* is the is the coefficient of the term for the *j*th pole *λ_j_* in the partial fraction expansion of *H*(*z*). The *λ* and *R* terms are computed from the eigenvalue decomposition of the companion form of the *A* parameter matrix. *λ* denotes the eigenvalues (and in this case, the roots of the denominator of *H*(*z*)) and *R* the outer product of the eigenvectors for a given mode.

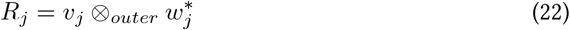

*R_j_* is a *m × m* matrix of rank one containing the coefficients for term *j* of the partial fraction expansion. This scales the mode into each node and connection in the system.

The number of terms in the expansion is the product of the number of channels in the dataset *m* and the MVAR model order *p*. The poles may either exist as single, real valued terms or in complex-conjugate pairs.

Through the identity,

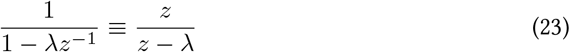

we can remove the powers on *z* and further simplify equation 21 to

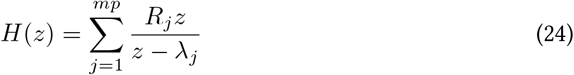

Equation 24 is also known as Gilbert’s realisation and it re-parametrises the system transfer function as a partial fraction expansion [30]. This realisation is valid when there are no repeated poles in *λ*. This approach has wide applications in the field of engineering controls systems and a modal analysis [57]. Equation 24 shows that *H*(*z*) from a linear system may be written entirely in terms of the roots of its time domain parameters. As such, the number of degree of freedom in the frequency domain representation of a linear system is completely determined by the number of roots of *a*. It follows that the configuration of *λ* in the *z*-plane forms a natural basis for a set of oscillatory modes as derived by [23]. Moreover, each mode (linked to a real valued or complex conjugate pair of pole) has a direct physical interpretation as an oscillator with a peak frequency and network structure throughout the system.

### 4.2 Computation of modal parameters

We can compute the polynomial roots *λ* of *H*(*z*) by finding the values at which the denominator polynomial of equation 14 evaluates to zero. These roots may be estimated by computing the eigenvalues of a matrix form of the polynomial. In the previous section, we discuss the denominator of *H* in terms of the 3-dimensional parameter matrix *A* however this does not permit an eigenvalue decomposition. Therefore we replace the order-p parameter matrix *A* with its companion form *C* as defined in equation 7.

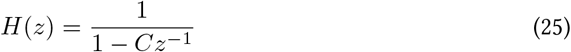

As *C* is a [*mp × mp*] square matrix, we can compute the eigenvalues decomposition with eigenvalues *λ* and eigenvectors *W*.

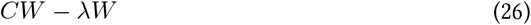

The eigenvalues are then the roots of its characteristic polynomial of *C* [58, section 7.1.1].

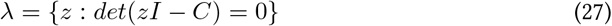

The roots are the set of values *λ* for which equation 27 evaluates to zero. This method for computing the roots of a polynomial from the eigenvalues of the companion matrix is commonly used in software (see documentation for *roots* in MATLAB or *numpy.roots* in Python’s Numpy library). Crucially, the characteristic polynomials of *A* and *C* are equivalent so their roots will describe the same dynamics. The construction of the square matrix *C* is convenient to permit this eigenvalue decomposition. There are many analytic and numerical approaches for computing the roots of univariate polynomials, but to our knowledge this is the most robust approach for finding the roots of a multivariate polynomial such as *A* where *m >* 1.

The eigenvectors associated with each eigenvalue are then the non-zero vectors for which equation 26 holds true.

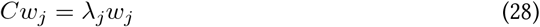

We are only interested in non-trivial solutions in which *w_j_* = 0 that occur when the determinant in equation 27 is equal to zero. The eigenvectors above are known as the right eigenvectors, the left eigenvectors can be computed as the the inverse of the conjugate transpose of the right eigenvectors.

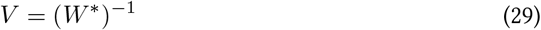

Once computed from the eigenvalue decomposition of *C*, the parameters *λ*, *W* and *V* can be plugged into the equations in sections 2.1 and 4.1 to define the spatio-spectral eigenmodes.

### 4.3 Software

All simulation, MVAR modelling and model decomposition steps are computed in Python 3.7.3 the Spectral Analysis In Linear Systems toolbox [59], https://vcs.ynic.york.ac.uk/analysis/sails and https://sails.readthedocs.io). MEG data pre-processing and beamforming was performed using Fieldtrip and the OHBA Software Library (https://github.com/OHBA-analysis/osl-core; [60]) in Matlab version R2019a on a cluster of x86-64 systems. The Bayesian statistical analysis was carried out in R version 3.5.2 using the *BRMS* (version 2.11.1; [61, 62]) and *loo* (version 2.2.0; [63]) packages. Full scripts for the preprocessing, data analysis and statistical assessment in simulated and HCP MEG data are available online (https://vcs.ynic.york.ac.uk/analysis/rs-mvar). The scripts include a tool for checking out the correct versions of the external toolboxes which are used.

### 4.4 Simulations

The MVAR Modal Decomposition is first explored with simulations. 20 realisations (representative of 20 participants) of 300 seconds of data from a 10 node network are generated with a sampling frequency of 128Hz. Each dataset is built from two subnetworks with different spatial and spectral profiles, the first is defined by a real-valued pole at 0Hz and the second by a complex-conjugate pair of poles between 8Hz and 12Hz, jittered across participants. Oscillatory data for these networks are generated by placing the poles within the z-plane and transforming them back to their polynomial form. These polynomials are then used as coefficients to filter white noise to produce oscillatory time-series. Each of the two oscillations are then projected through the network using a set of weights defining the relative strength of the oscillation in each of the 10 nodes. Finally the two oscillatory networks are added together with white noise to create the final signal.

Each network is described with an order 5 MVAR model. The Fourier-based cross and power spectral density (CPSD) *P_f_* is computed and the averaged within two frequency bands of interest 0–4Hz and 8–12Hz reflecting the two simulated oscillations. The modal decomposition is then computed and a modal form of *P_m_* split into three reduced models, two models for the poles which survive the permutation thresholding procedure and a residual model. The poles-of-interest for the simulation are taken as those which are identified as surviving the thresholding procedure. The surviving poles are then assigned to the low or high frequency band of interest based on having a characteristic frequency lying within 4Hz of the relevant frequency (the same ranges as used for the Fourier analysis).

### 4.5 Resting state MEG data

Resting-state MEG datasets from 79 participants in the Human Connectome Project [24, 25] were used. Each participant underwent three separate runs of a 6-minute eyes-open resting state protocol MEG data were collected using a 4D Neuroimaging WH-3600 scanner, equipped with 248 magnetometer sensor channels and 23 reference channels, and were sampled at 2034.51Hz. Participant headshapes were digitised using a polhemus tracker system prior to MEG data collection.

The HCP pre-processed resting state MEG datasets were used along with the room noise recordings for the relevant session and information regarding the ICA components from the de-noising process. Co-registrations for the MEG and MRI data for each participant were taken from the models provided by the HCP project.

Seventy-eight areas from the AAL2 atlas [27, 64] were used as target regions of interest. Beamformer weights were calculated for locations on an 8mm-spaced grid spaced inside each of the regions of interest. A Linearly-Constrained Minimum Variance beamformer [26, 65] at the orientation which showed maximum power. The source virtual electrode time-series were then resampled to 240Hz. The individual time-series from the grid locations within each region were then reduced to a single time-series per region by taking the first principal component across the voxels within the region. The time-series across all regions were then orthogonalised to reduce the impact of spatial leakage [36]. Finally, the beamformed time-series were downsampled 2-fold using a windowed fourier-domain method, giving a final sampling rate of 120Hz.

### 4.6 Model order selection

Prior to analysis of an autoregressive model of any dataset, the model order *p* must be selected. This choice can be informed by metrics such as Akaike’s Information Criterion (AIC: [66], however this often gives a monotonically decreasing profile with no clear optimal model [67–70]. Even if there is a local minimum in the AIC time-domain metric, this does not guarantee that the resulting power spectrum will provide a good representation of the data. Therefore, we choose a model order which both produces a good power spectrum and has a low AIC.

### 4.7 Model fitting and validation

MVAR Models were fitted with order 12 across all 78 parcels in the HCP data. 12 was chosen by a combination of the AIC and manual inspection of the model spectra. An order of 12 produced good spectra and was not before an inflection point in the AIC.

After fitting, the models were checked for stability (using the Stability Index (SI): [71, pages 15–16]), residual autocorrelation (using the Durbin-Watson index: [72]) and variance explained. The models were able to fit between 21–29% of variance (mean=24.990%, SD=4.003%) within each recording session. We consider this to be a good proportion of variance to explain with a single stationary and linear model of a whole brain functional parcellation. All models were stable, having SI values below 1 (mean=.959, SD=.016) and no substantial autocorrelation could be found in the residuals according to the Durbin-Watson test (mean=2.003, SD*<*.001).

Once the MVAR models (*A* matrix) were fitted for each scan session, the transfer function *H* and spectral matrix *S* were computed between 0 and 48Hz using the Fourier method. The *S* matrices were averaged within the set of specified frequency bands to summarise the frequency-specific spatial topologies captured by the MVAR models.

### 4.8 Fourier and SSE network connectivity estimation

We validate that a single MVAR model is able to describe the spatial and spectral content of a whole brain functional connectome estimated from MEG data using a standard Fourier-based approach. The system transfer function is estimated using the Fourier equation 2 for all frequencies between 0-60Hz in 100 steps. Subsequently the spectral matrix is computed for each frequency using the *H*(*f*) and the residual covariance matrix Σ. Finally, we integrate within a set of pre-specified frequency bands to summarise how the network structure of oscillatory brain networks changes across frequency.

### 4.9 Modal decomposition and non-parametric permutation

The modal decomposition of each MVAR model was computed using the methods described above and the peak frequency, damping time, *H* and *S* were computed for each mode. The modal decomposition of a system returns *m * p* modes which could number in the hundreds or thousand for a large system. Many of these modes are likely to be modelling noisy characteristics of the system or its measurement rather than physiologically interesting oscillatory activity. In order to select the most dynamically relevant modes, a non-parametric permutation testing method was used on the damping times of the modes. Each individual timeseries was split into non-overlapping temporal epochs resulting in a 3d data array [channels x samples x epochs]. Permutations are carried out by randomising both the channels and epochs in order to construct null datasets in which the relationships between nodes have been destroyed whilst maintaining the overall spectral nature of the data. At each permutation, a MVAR model is estimated on the surrogate dataset and the modal decomposition computed. A maximum statistic method was then used [73] in which the maximum damping time of all of the modes within the given model was entered into the null distribution. This was repeated for each permutation, resulting in a null distribution of damping times for each participant, for each run. A threshold which represented the 1% tail of the null distribution was then established in this way for each run, for each participant. Modes were then selected from the un-permuted data using these individual damping time thresholds.

### 4.10 Spatial Principal Components Analysis of SSE networks

Patterns of spatial and network variation in the SSE surviving the permutation scheme was performed using a principal components analysis (PCA). The [nnodes x nnodes] PSD matrices for the significant SSEs were vectorised and concatenated a [modes x nnodes*nnodes] matrix and demeaned before a PCA was used to identify the principal axes of variation across the connections within the network across modes. The components of the PCA then show patterns in the spatial distribution of oscillatory power across a number of modes regardless of the characteristic frequency of the modes which significantly contribute to the component. Whilst each network is computed at its peak resonant frequency, these resonances are free to vary (within the specified alpha range) across networks both between and within individuals.

The PCA was computed for subsets of SSE whose peak frequency lies within each of three frequency bands. Theta (1–7Hz), alpha (7–13Hz) and beta (13–30Hz). Crucially, the inclusion of an SSE in a band depends only on its peak frequency. Once included, all information on the network structure within that SSE is included in the analysis, even if part of the spectral peak goes outside the specified band. Reproducibility of the components arising from the PCA were assessed using a split-half correlation. 500 split halves of the SSEs included in a PCA were computed and the PCA computed on each half independently before the spatial components of each half are then correlated and stored. Both the split-half correlation and proportion of variance explained by each component was used in determining whether the component was included in further analyses. The components describe the pattern of variability across space captured by that PC whilst the PC-score indicates the extent to which that shape is expressed in each individual SSE. An example spatial map is computed for maximum and minimum observed score in each PC by projecting that score back into the original data-space.

Relatively few SSEs survived the permutation scheme in the theta and beta bands.

Potentially as a result, the PCA components from these bands also showed relatively low reproducibility. For completeness, the SSE and the first 4 PCs for these bands are included in section 9.5. In contrast, over 1,200 SSEs were included in the alpha band and the first two PCA components showed high variance explained and split-half reliability. These are interpreted in the main text and carried forward for further statistical analysis.

### 4.11 Relationship between mode frequency and PC projection score

In order to examine whether there was a relationship between the frequency of each mode and the score with which it projected onto a given component, we performed a Bayesian linear regression using the BRMS package [61, 62]. For each PC, we scaled the scores by its standard deviation and fit a model of *Score_Scaled_ Frequency* + (1 | *Participant*), allowing an overall change in mean frequency per-participant. Model inference was performed using the standard NUTS sampler used by STAN through BRMS.

The prior for the Frequency parameter was chosen to be normally distributed with a mean of 0 and a standard deviation of 1; reflecting our default position that there was no a-prior reason to expect frequency to vary with score. Altering the standard deviation of the prior to other plausible ranges had no significant effect on the overall results. Examination of diagnostic plots showed that the parameters have converged in all cases.

To assess whether there was a relationship for each principal component, we fit an intercept-only model and a model with frequency as an additional linear regressor and compared evidence for the models using a Leave-One-Out (LOO) cross-validation methods [63]. Our criteria for determining that a model with the additional frequency regressor has more evidence is that the difference in LOO should be more than twice the estimate of the LOO standard standard error. For models where the model with frequency was assessed as having more evidence, we then report and assess the magnitude of the frequency parameter in the full model along with is 95% Credible Interval (CI). Due to the scaling of the scores, the frequency parameters is expressed in terms of the standard deviation of the score.

## 6 Acknowledgments

The authors would like to thank Sam Johnson and Catharina Zich. The HCP data were provided by the Human Connectome Project, WU-Minn Consortium (Principal Investigators: David Van Essen and Kamil Ugurbil; 1U54MH091657) funded by the 16 NIH Institutes and Centers that support the NIH Blueprint for Neuroscience Research; and by the McDonnell Center for Systems Neuroscience at Washington University.

## 7 Data & Code Availability

All scripts for data simulation, processing, analysis and visualisation in this paper are available online at https://vcs.ynic.york.ac.uk/analysis/rs-mvar-scripts. The HCP data used for the human MEG component of the paper is available from the Human Connectome Project.

MEG data from this study are available to download through the Human Connectome Project (HCP; www.humanconnectome.org). Prior to downloading data, users must register with the HCP and agreed to the data use terms (https://www.humanconnectome.org/study/hcp-young-adult/data-use-terms).

## 8 Author Contributions

**Conceptualization:** Andrew Qyinn, Gary Green, Mark Hymers. **Data curation:** Andrew Qyinn, Mark Hymers. **Formal Analysis:** Andrew Qyinn, Mark Hymers **Funding Acquisition:** Andrew Qyinn, Mark Hymers **Methodology:** Andrew Qyinn, Mark Hymers **Software:** Andrew Qyinn, Mark Hymers **Validation:** Andrew Qyinn, Mark Hymers **Visualisation:** Andrew Qyinn, Mark Hymers **Writing - original draft:** Andrew Qyinn, Mark Hymers **Writing - review & editing** : Andrew Qyinn, Gary Green, Mark Hymers

## 9 Supporting information

### 9.1 Notation

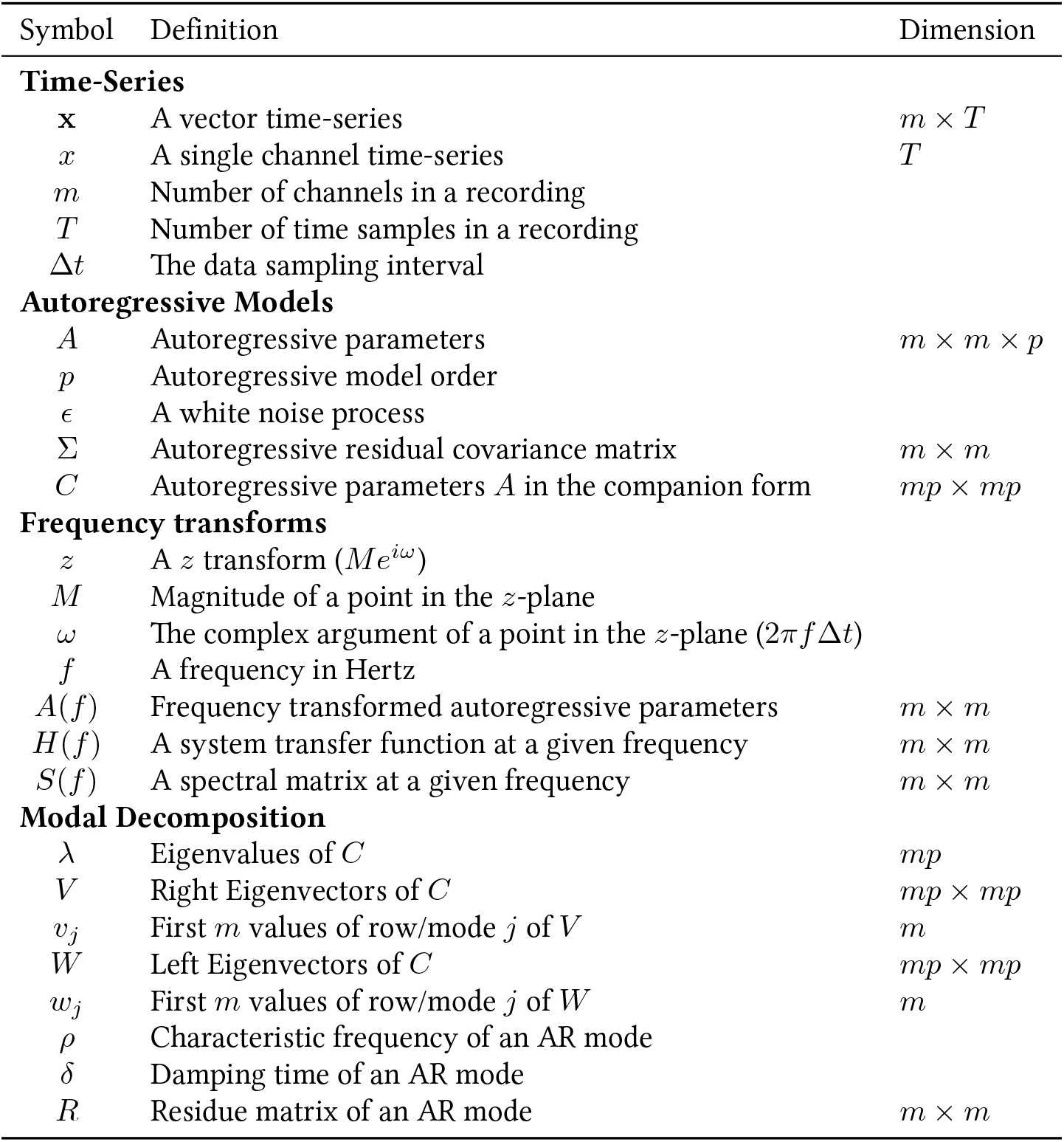

### 9.2 Individual realisation results for group simulation

In Figure S.1 we present the results for each of the 20 individuals in the simulation. These results are analagous to those shown for the example participant in panel F of Figure 1.

**Fig S.1.**
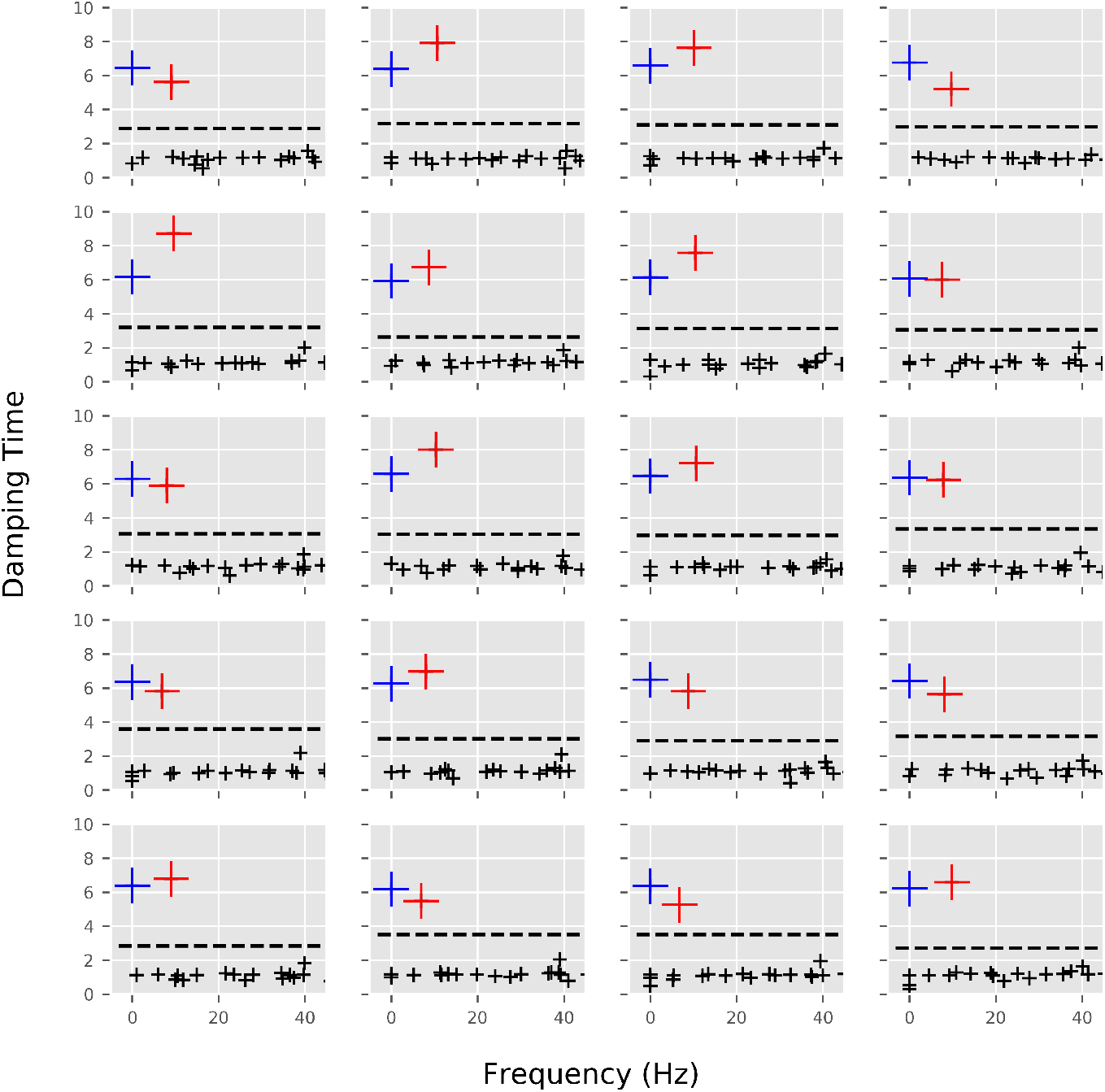
Replication of figure 1F for all 20 individual simulation realisations. Damping-time of each mode as a function of frequency. The slope (blue cross) and 9Hz peak (red crosses) have the longest damping times indicating that the dynamics in these modes are relatively un-damped and possibly of greater dynamical importance to the system. The dashed line indicates the 99% significance threshold computed for each individual run via the permutation scheme described in the main text.

### 9.3 Model Order Comparisons

Figure S.2 shows a comparison of MVAR models across orders from 2 to 16 in steps of two. figure S.3 shows the comparable data for distribution of thresholded modes as computed in the manner discussed in the main text.

### 9.4 Split-half reliability of alpha components

In Figure S.4 we show the split-half reliabilities of the 50 principal components of the alpha decomposition over 500 iterations.

**Fig S.2.**
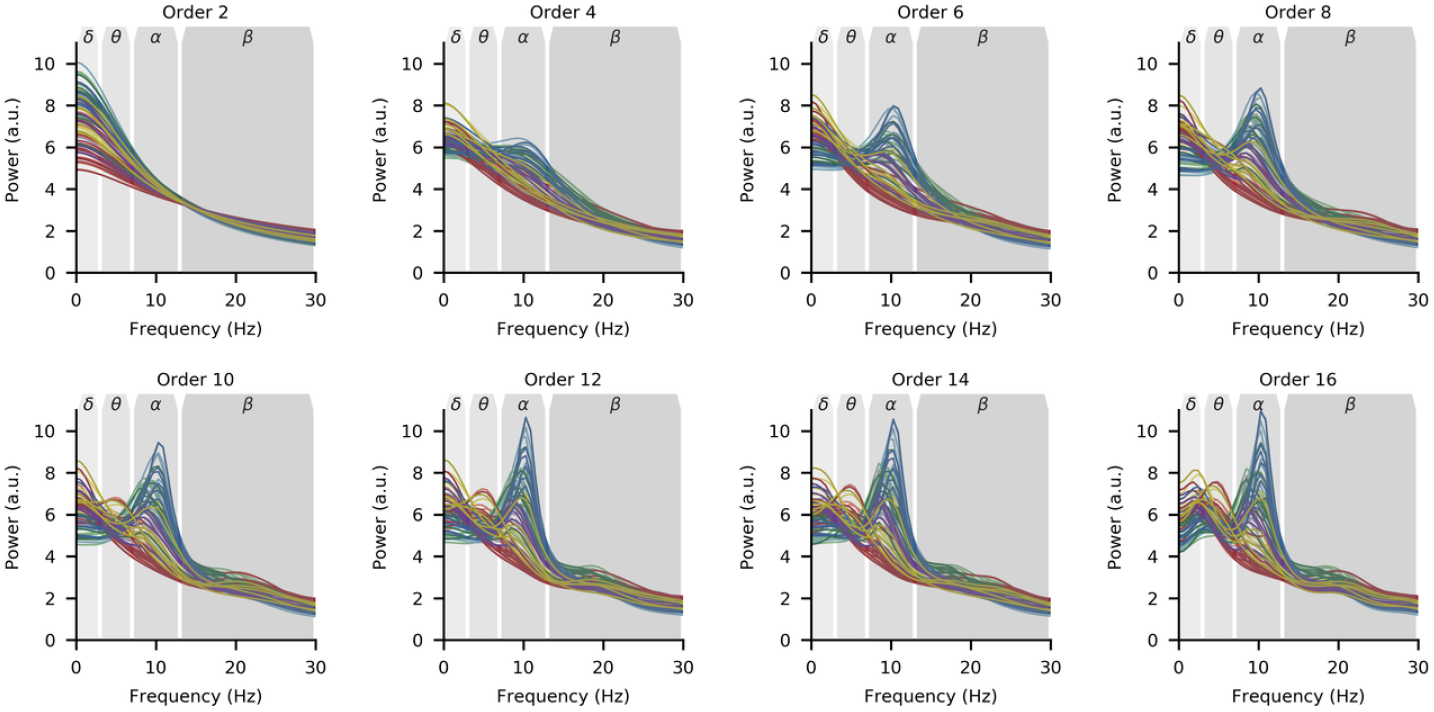
Spectrum estimation from MVAR models computed at orders 2 through 16 in steps of 2.

**Fig S.3.**
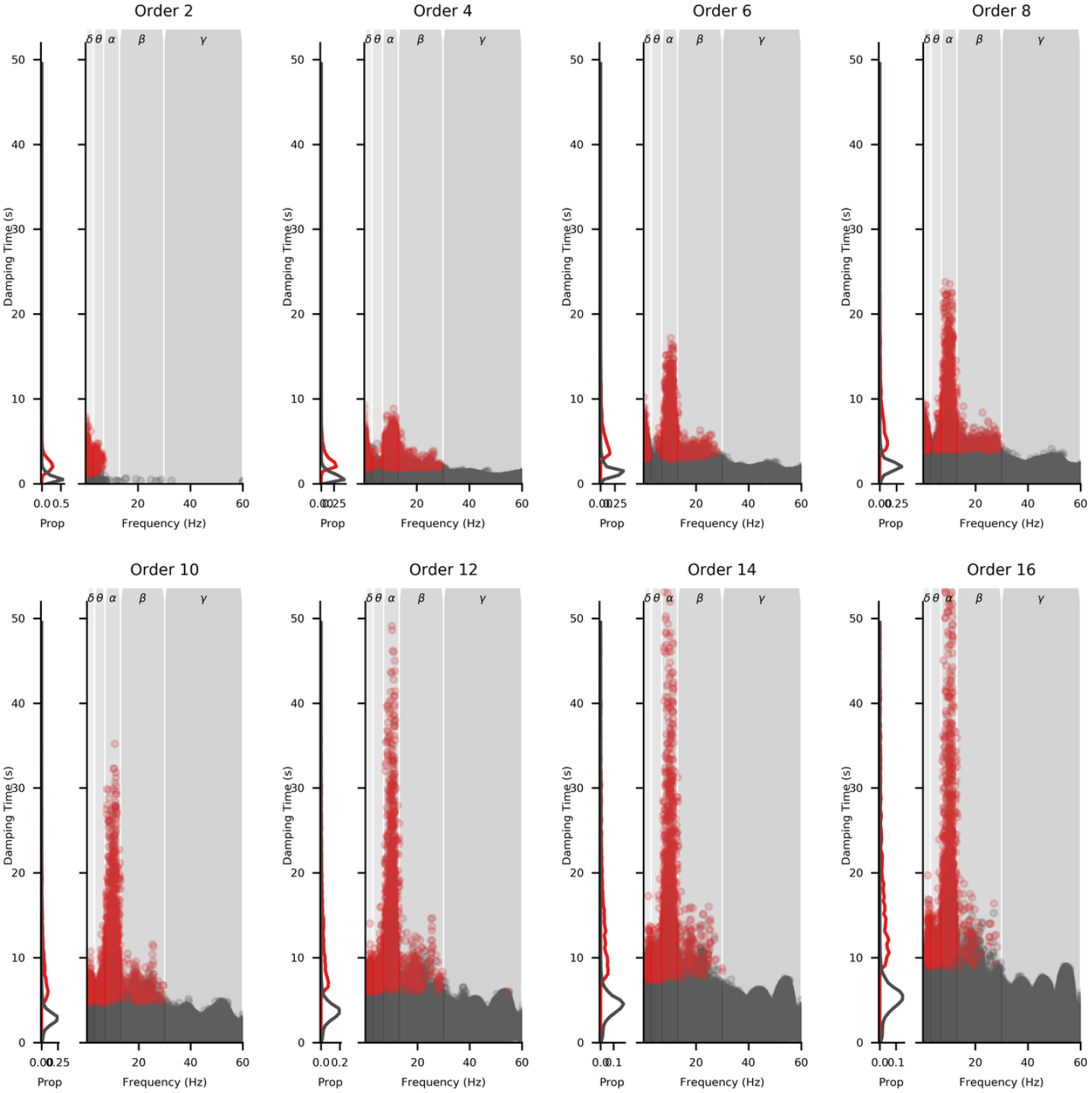
Surviving mode distributions computed from MVAR models of orders 2 through 16 in steps of 2.

**Fig S.4.**
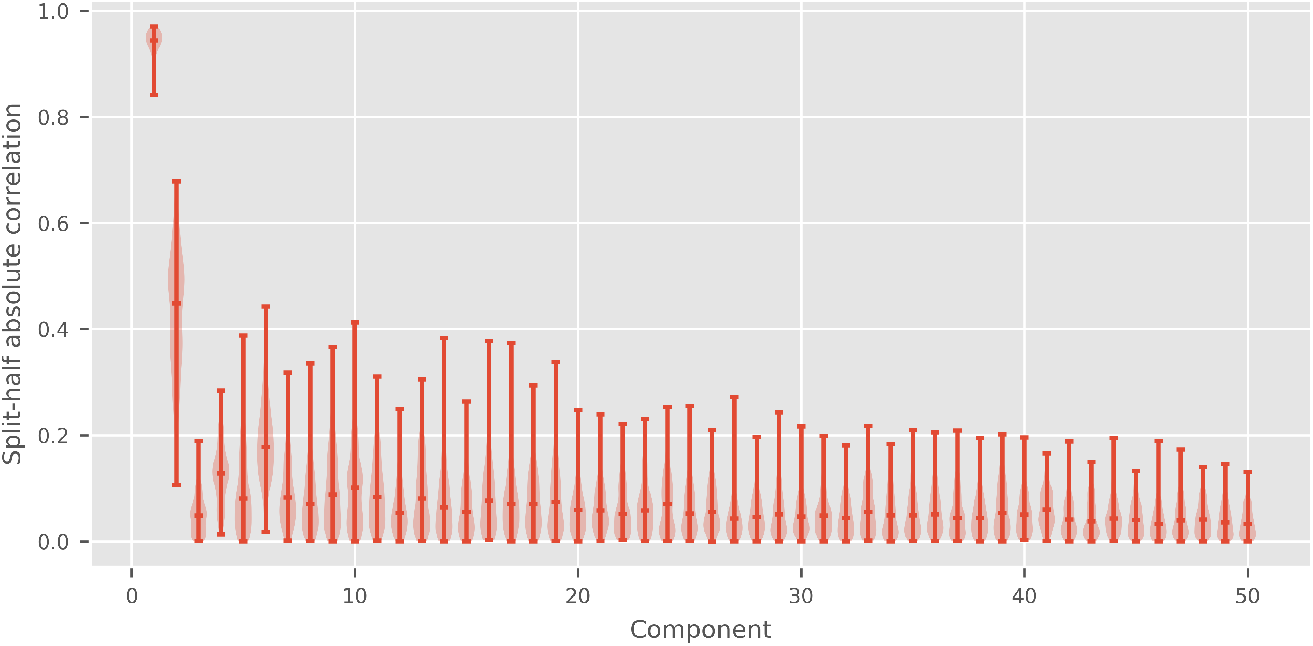
Split-half reliability of the 50 principal components for the alpha SSE decomposition. The split-half correlation values are shown as the distributions of the the absolute value of the correlations (as the sign of the eigenvector-based map is confounded with the sign of the eigenvalue). The horizontal bar on each plot shows the median value.

### 9.5 Spatio-Spectral Eigenmodes in the theta and beta bands

In this section, we show figures for the theta and beta band data which are analagous to those shown for the alpha band data.

Figures S.5 and S.6 show the split-half data (analaogus to the data reported in figure S.4 for the alpha band) whilst figure S.7 shows the surface power for the first four components of the theta and beta bands; this can be compared with figure 8 for the first two components of the alpha band analysis.

**Fig S.5.**
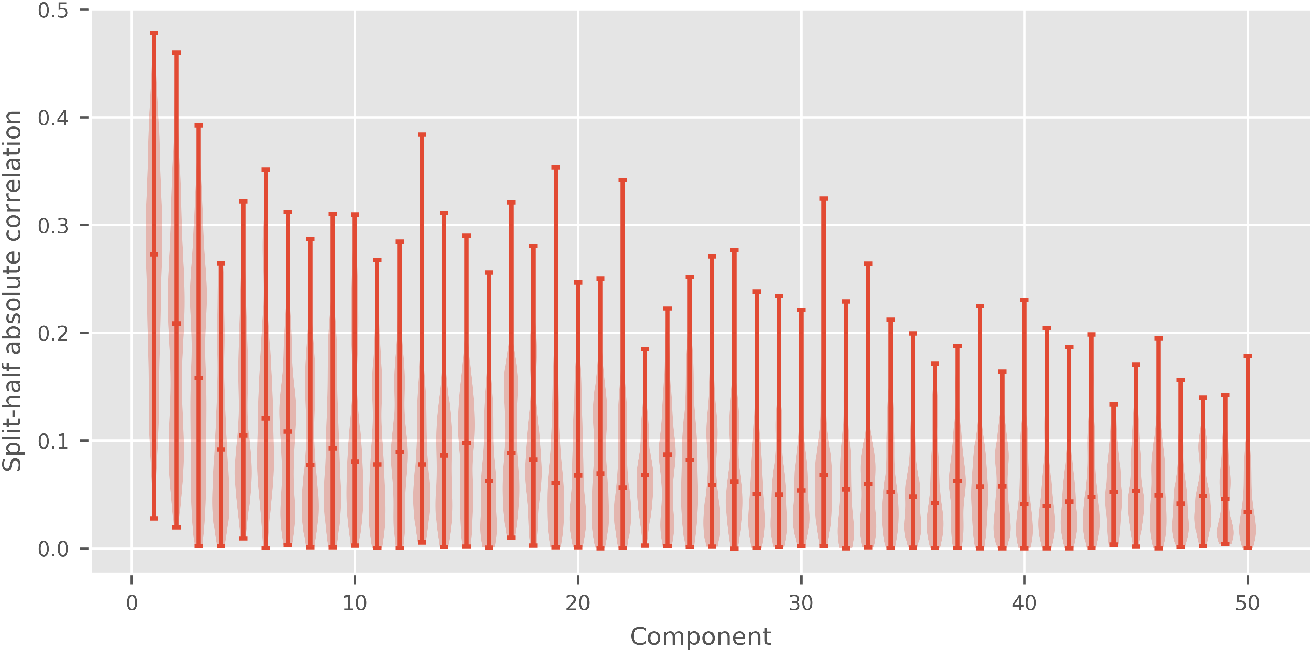
Split-half reliability of the 50 principal components for the theta SSE decomposition. The split-half correlation values are shown as the distributions of the the absolute value of the correlations (as the sign of the eigenvector-based map is confounded with the sign of the eigenvalue). The horizontal bar on each plot shows the median value.

### 9.6 Relationship between PC-score and frequency for the alpha band

**Fig S.6.**
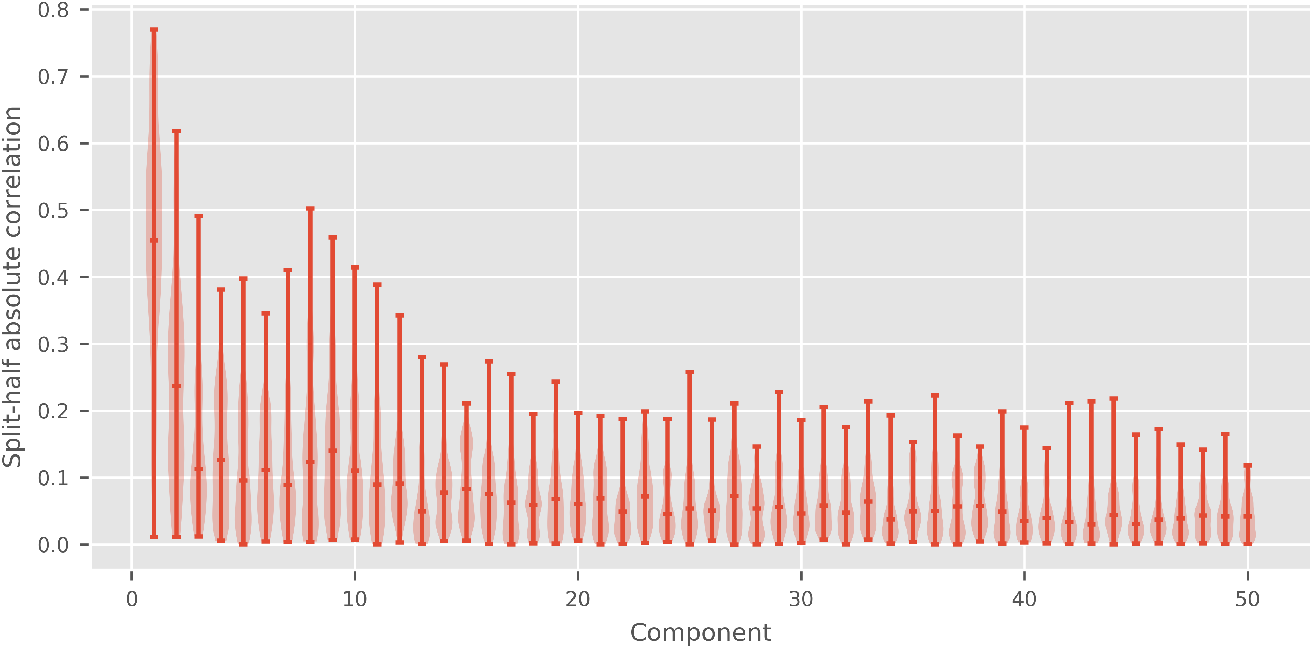
Split-half reliability of the 50 principal components for the beta SSE decomposition. The split-half correlation values are shown as the distributions of the the absolute value of the correlations (as the sign of the eigenvector-based map is confounded with the sign of the eigenvalue). The horizontal bar on each plot shows the median value.

**Fig S.7.**
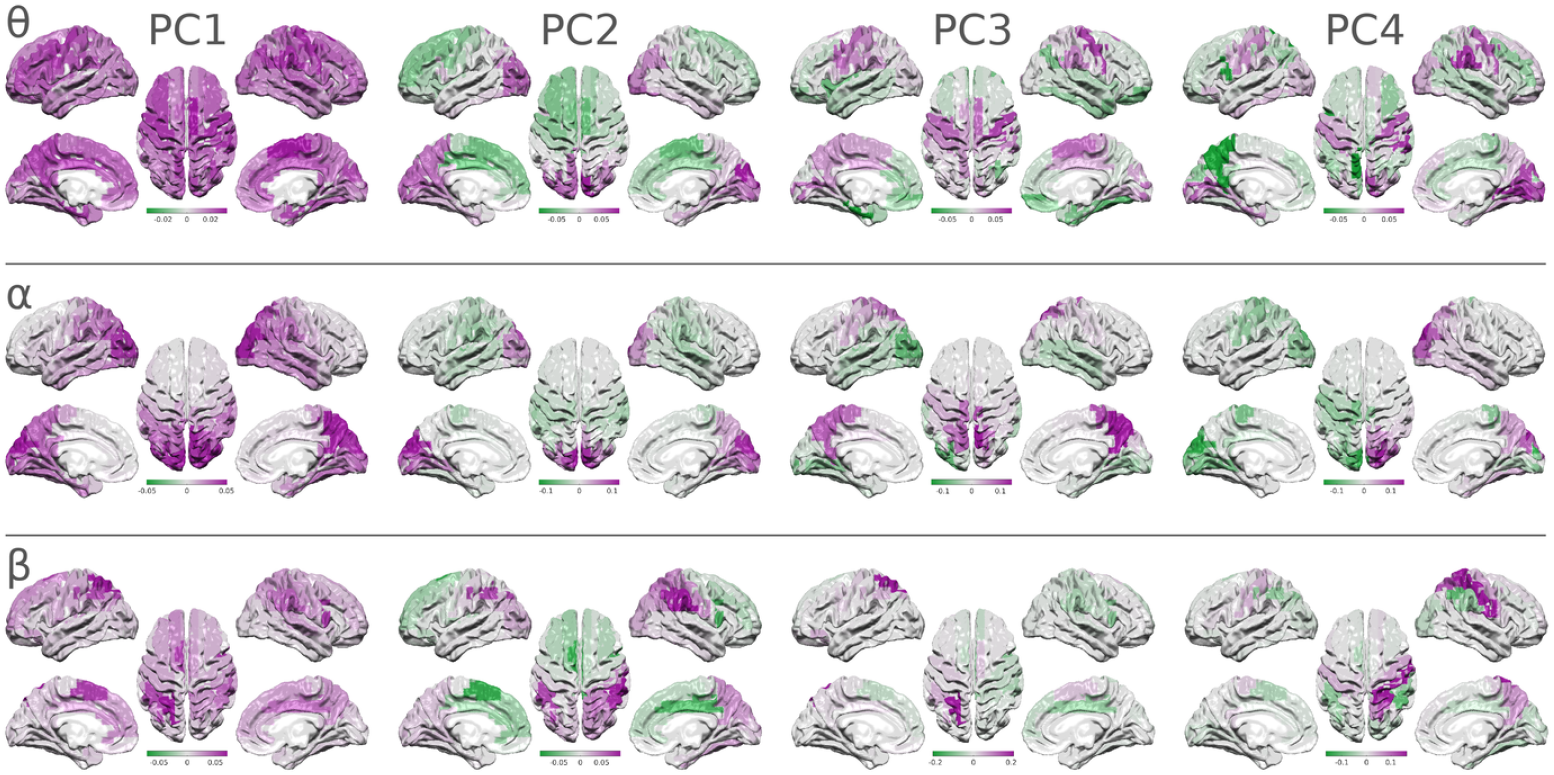
Surface visualisations of the first four principal components for each of the theta, alpha and beta bands. These can be compared with the first two alpha band components shown in figure 8 of the main paper.

**Fig S.8.**
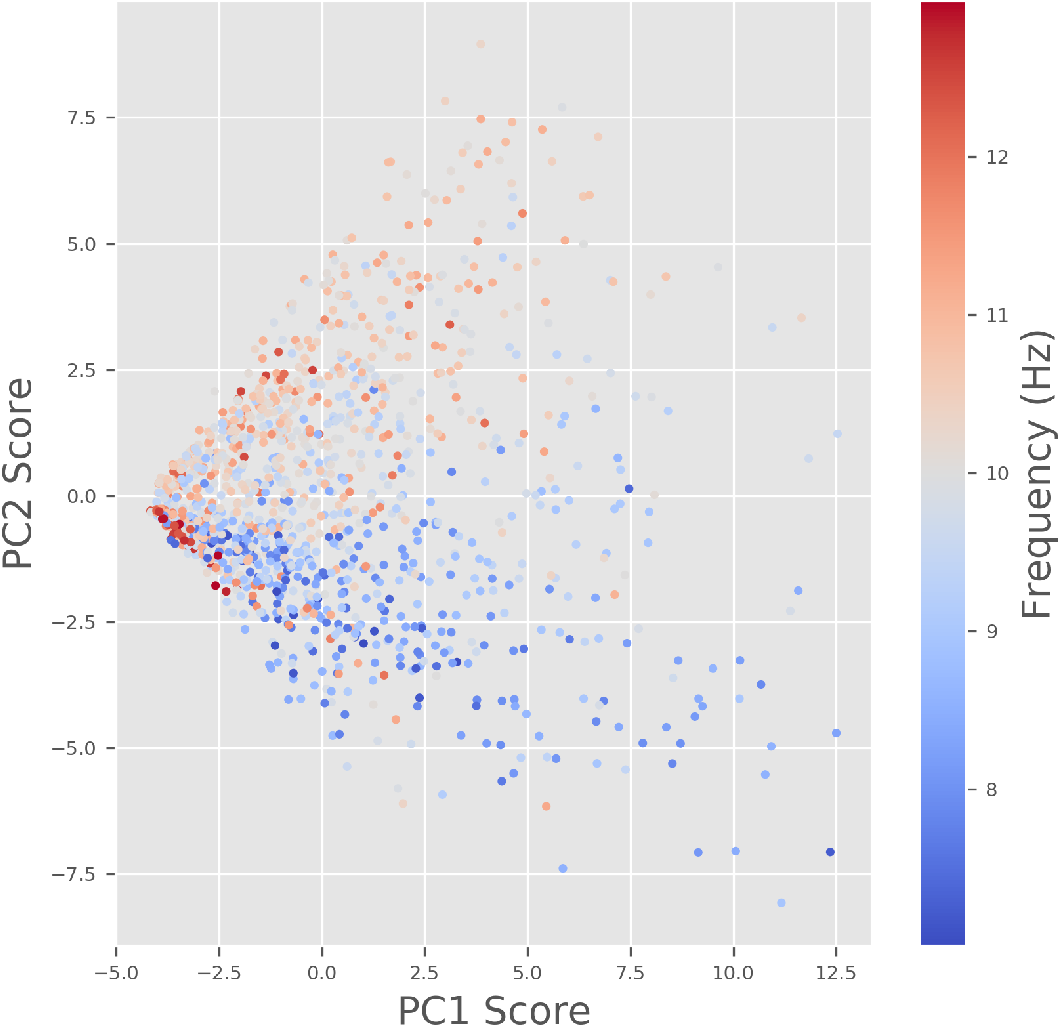
Scatter plot of the PC-scores for the first two components of the results in the alpha band. Each point is a single SSE and colour indicates peak frequency. The x-axis contains the scores for PC-1 which correspond to overall alpha power. The PC scores are orthogonal and have no linear correlation. Low PC-1 scores indicate SSEs with low overall power whilst high scores indicate SSEs with strong alpha networks which, in turn, have greater variability in PC-2. The frequency correlation with PC2 can be seen as a greater density of high frequencies (red colours) above zero in the y-axis.

## References

1. Bressler SL. Large-scale cortical networks and cognition. Brain Research Reviews. 1995;20(3):288–304. doi:10.1016/0165-0173(94)00016-i.

2. Fries P. A mechanism for cognitive dynamics: neuronal communication through neuronal coherence. Trends in Cognitive Sciences. 2005;9(10):474–480. doi:10.1016/j.tics.2005.08.011.

3. Berger H. Über das elektrenkephalogramm des menschen. European archives of psychiatry and clinical neuroscience. 1929;87(1):527–570.

4. da Silva FL. Neural mechanisms underlying brain waves: from neural membranes to networks. Electroencephalography and Clinical Neurophysiology. 1991;79(2):81–93. doi:10.1016/0013-4694(91)90044-5.

5. Klimesch W, Sauseng P, Hanslmayr S. EEG alpha oscillations: The inhibition–timing hypothesis. Brain Research Reviews. 2007;53(1):63–88. doi:10.1016/j.brainresrev.2006.06.003.

6. Jensen O, Mazaheri A. Shaping Functional Architecture by Oscillatory Alpha Activity: Gating by Inhibition. Frontiers in Human Neuroscience. 2010;4. doi:10.3389/fnhum.2010.00186.

7. Klimesch W. EEG alpha and theta oscillations reflect cognitive and memory performance: a review and analysis. Brain Research Reviews. 1999;29(2-3):169–195. doi:10.1016/s0165-0173(98)00056-3.

8. Haegens S, Cousijn H, Wallis G, Harrison PJ, Nobre AC. Inter- and intra-individual variability in alpha peak frequency. NeuroImage. 2014;92:46–55. doi:10.1016/j.neuroimage.2014.01.049.

9. Osipova D, Ahveninen J, Jensen O, Ylikoski A, Pekkonen E. Altered generation of spontaneous oscillations in Alzheimer’s disease. NeuroImage. 2005;27(4):835–841. doi:10.1016/j.neuroimage.2005.05.011.

10. Poza J, Hornero R, Abásolo D, Fernández A, García M. Extraction of spectral based measures from MEG background oscillations in Alzheimer’s disease. Medical Engineering & Physics. 2007;29(10):1073–1083. doi:10.1016/j.medengphy.2006.11.006.

11. Hughes LE, Henson RN, Pereda E, Brunña R, López-Sanz D, Qyinn AJ, et al. Biomagnetic biomarkers for dementia: A pilot multicentre study with a recommended methodological framework for magnetoencephalography. Alzheimer’s & Dementia: Diagnosis, Assessment & Disease Monitoring. 2019;11(1):450–462. doi:10.1016/j.dadm.2019.04.009.

12. Garcés P, Vicente R, Wibral M, Pineda-Pardo JÁ, López ME, Aurtenetxe S, et al. Brain-wide slowing of spontaneous alpha rhythms in mild cognitive impairment. Frontiers in Aging Neuroscience. 2013;5. doi:10.3389/fnagi.2013.00100.

13. López-Sanz D, Bruña R, Garcés P, Camara C, Serrano N, Rodríguez-Rojo IC, et al. Alpha band disruption in the AD-continuum starts in the Subjective Cognitive Decline stage: a MEG study. Scientific Reports. 2016;6(1). doi:10.1038/srep37685.

14. Engels MMA, Hillebrand A, van der Flier WM, Stam CJ, Scheltens P, van Straaten ECW. Slowing of Hippocampal Activity Correlates with Cognitive Decline in Early Onset Alzheimer’s Disease. An MEG Study with Virtual Electrodes. Frontiers in Human Neuroscience. 2016;10. doi:10.3389/fnhum.2016.00238.

15. Peraza LR, Cromarty R, Kobeleva X, Firbank MJ, Killen A, Graziadio S, et al. Electroencephalographic derived network differences in Lewy body dementia compared to Alzheimer’s disease patients. Scientific Reports. 2018;8(1). doi:10.1038/s41598-018-22984-5.

16. Hari R. Human cortical oscillations: a neuromagnetic view through the skull. Trends in Neurosciences. 1997;20(1):44–49. doi:10.1016/s0166-2236(96)10065-5.

17. Ciulla C, Takeda T, Endo H. MEG Characterization of Spontaneous Alpha Rhythm in the Human Brain. Brain Topography. 1999;11(3):211–222. doi:10.1023/a:1022233828999.

18. Wens V, Bourguignon M, Goldman S, Marty B, de Beeck MO, Clumeck C, et al. Inter- and Intra-Subject Variability of Neuromagnetic Resting State Networks. Brain Topography. 2014;27(5):620–634. doi:10.1007/s10548-014-0364-8.

19. Colclough GL, Smith SM, Nichols TE, Winkler AM, Sotiropoulos SN, Glasser MF, et al. The heritability of multi-modal connectivity in human brain activity. eLife. 2017;6:e20178. doi:10.7554/eLife.20178.

20. Sokoliuk R, Mayhew SD, Aquino KM, Wilson R, Brookes MJ, Francis ST, et al. Two Spatially Distinct Posterior Alpha Sources Fulfill Different Functional Roles in Attention. The Journal of Neuroscience. 2019;39(36):7183–7194. doi:10.1523/jneurosci.1993-18.2019.

21. Barzegaran E, Vildavski VY, Knyazeva MG. Fine Structure of Posterior Alpha Rhythm in Human EEG: Frequency Components, Their Cortical Sources, and Temporal Behavior. Scientific Reports. 2017;7(1). doi:10.1038/s41598-017-08421-z.

22. von Storch H, Bürger G, Schnur R, von Storch JS. Principal Oscillation Patterns: A Review. Journal of Climate. 1995;8(3):377–400. doi:10.1175/1520-0442(1995)008¡0377:popar¿2.0.co;2.

23. Neumaier A, Schneider T. Estimation of parameters and eigenmodes of multivariate autoregressive models. ACM Transactions on Mathematical Software. 2001;27(1):27–57. doi:10.1145/382043.382304.

24. Larson-Prior LJ, Oostenveld R, Della Penna S, Michalareas G, Prior F, Babajani-Feremi A, et al. Adding dynamics to the Human Connectome Project with MEG. Neuroimage. 2013;80:190–201.

25. Van Essen DC, Smith SM, Barch DM, Behrens TE, Yacoub E, Ugurbil K, et al. The WU-Minn Human Connectome Project: an overview. Neuroimage. 2013;80:62–79.

26. Van Veen BD, Van Drongelen W, Yuchtman M, Suzuki A. Localization of brain electrical activity via linearly constrained minimum variance spatial filtering. IEEE Trans Biomed Eng. 1997;44(9):867–880. doi:10.1109/10.623056.

27. Rolls ET, Joliot M, Tzourio-Mazoyer N. Implementation of a new parcellation of the orbitofrontal cortex in the automated anatomical labeling atlas. NeuroImage. 2015;122:1–5. doi:10.1016/j.neuroimage.2015.07.075.

28. Blinowska KJ. Review of the methods of determination of directed connectivity from multichannel data. Medical & Biological Engineering & Computing. 2011;49(5):521–529. doi:10.1007/s11517-011-0739-x.

29. Baccalá LA, Sameshima K. Partial directed coherence: a new concept in neural structure determination. Biological Cybernetics. 2001;84(6):463–474. doi:10.1007/pl00007990.

30. Gilbert EG. Controllability and observability in multivariable control systems. Journal of the Society for Industrial and Applied Mathematics, Series A: Control. 1963;1(2):128–151.

31. Kailath T. Linear systems. Prentice-Hall; 1980.

32. Brookes MJ, Woolrich M, Luckhoo H, Price D, Hale JR, Stephenson MC, et al. Investigating the electrophysiological basis of resting state networks using magnetoencephalography. Proceedings of the National Academy of Sciences. 2011;108(40):16783–16788. doi:10.1073/pnas.1112685108.

33. Hipp JF, Hawellek DJ, Corbetta M, Siegel M, Engel AK. Large-scale cortical correlation structure of spontaneous oscillatory activity. Nature neuroscience. 2012;15(6). doi:10.1038/nn.3101.

34. Marzetti L, Penna SD, Snyder AZ, Pizzella V, Nolte G, de Pasquale F, et al. Frequency specific interactions of MEG resting state activity within and across brain networks as revealed by the multivariate interaction measure. NeuroImage. 2013;79:172–183. doi:10.1016/j.neuroimage.2013.04.062.

35. Wens V, Mary A, Bourguignon M, Goldman S, Marty B, beeck MOd, et al. About the electrophysiological basis of resting state networks. Clinical Neurophysiology. 2014;125(8):1711–1713. doi:10.1016/j.clinph.2013.11.039.

36. Colclough GL, Brookes MJ, Smith SM, Woolrich MW. A symmetric multivariate leakage correction for MEG connectomes. NeuroImage. 2015;117:439–448. doi:10.1016/j.neuroimage.2015.03.071.

37. Hillebrand A, Barnes GR, Bosboom JL, Berendse HW, Stam CJ. Frequency-dependent functional connectivity within resting-state networks: An atlas-based MEG beamformer solution. NeuroImage. 2012;59(4):3909–3921. doi:10.1016/j.neuroimage.2011.11.005.

38. Qyinn AJ, van Ede F, Brookes MJ, Heideman SG, Nowak M, Seedat ZA, et al. Unpacking Transient Event Dynamics in Electrophysiological Power Spectra. Brain Topography. 2019;32:1020–1034. doi:10.1007/s10548-019-00745-5.

39. Clayton MS, Yeung N, Kadosh RC. The many characters of visual alpha oscillations. European Journal of Neuroscience. 2017;48(7):2498–2508. doi:10.1111/ejn.13747.

40. Jensen O, Bonnefond M, VanRullen R. An oscillatory mechanism for prioritizing salient unattended stimuli. Trends in Cognitive Sciences. 2012;16(4):200–206. doi:10.1016/j.tics.2012.03.002.

41. van Dijk H, Schoffelen JM, Oostenveld R, Jensen O. Prestimulus Oscillatory Activity in the Alpha Band Predicts Visual Discrimination Ability. Journal of Neuroscience. 2008;28(8):1816–1823. doi:10.1523/jneurosci.1853-07.2008.

42. Huntenburg JM, Bazin PL, Margulies DS. Large-Scale Gradients in Human Cortical Organization. Trends in Cognitive Sciences. 2018;22(1):21–31. doi:10.1016/j.tics.2017.11.002.

43. Zhang H, Watrous AJ, Patel A, Jacobs J. Theta and Alpha Oscillations Are Traveling Waves in the Human Neocortex. Neuron. 2018;98(6):1269–1281.e4. doi:10.1016/j.neuron.2018.05.019.

44. Pineda JA. The functional significance of mu rhythms: Translating “seeing” and “hearing” into “doing”. Brain Research Reviews. 2005;50(1):57–68. doi:10.1016/j.brainresrev.2005.04.005.

45. Qyirk M, Liu B. Improving resolution for autoregressive spectral estimation by decimation. IEEE Transactions on Acoustics, Speech, and Signal Processing. 1983;31(3):630–637. doi:10.1109/tassp.1983.1164124.

46. Gersch W, Yonemoto J. Parametric time series models for multivariate EEG analysis. Computers and Biomedical Research. 1977;10(2):113–125. doi:10.1016/0010-4809(77)90029-5.

47. Franaszczuk PJ, Blinowska KJ. Linear model of brain electrical activity – EEG as a superposition of damped oscillatory modes. Biological Cybernetics. 1985;53(1):19–25. doi:10.1007/bf00355687.

48. Wright JJ, Kydd RR, Sergejew AA. Autoregression models of EEG. Biological Cybernetics. 1990;62(3):201–210. doi:10.1007/bf00198095.

49. Mullen T, Worrell G, Makeig S. Multivariate principal oscillation pattern analysis of ICA sources during seizure. In: 2012 Annual International Conference of the IEEE Engineering in Medicine and Biology Society. IEEE; 2012.Available from: https://doi.org/10.1109/embc.2012.6346575.

50. Juang JN, Pappa RS. An eigensystem realization algorithm for modal parameter identification and model reduction. Journal of Guidance, Control, and Dynamics. 1985;8(5):620–627. doi:10.2514/3.20031.

51. Hunyadi B, Camps D, Sorber L, Paesschen WV, Vos MD, Huffel SV, et al. Block term decomposition for modelling epileptic seizures. EURASIP Journal on Advances in Signal Processing. 2014;2014(1). doi:10.1186/1687-6180-2014-139.

52. Schmid PJ. Dynamic mode decomposition of numerical and experimental data. Journal of Fluid Mechanics. 2010;656:5–28. doi:10.1017/s0022112010001217.

53. Casorso J, Kong X, Chi W, Ville DVD, Yeo BTT, Liégeois R. Dynamic mode decomposition of resting-state and task fMRI. NeuroImage. 2019;194:42–54. doi:10.1016/j.neuroimage.2019.03.019.

54. Brunton BW, Johnson LA, Ojemann JG, Kutz JN. Extracting spatial–temporal coherent patterns in large-scale neural recordings using dynamic mode decomposition. Journal of Neuroscience Methods. 2016;258:1–15. doi:10.1016/j.jneumeth.2015.10.010.

55. Shiraishi Y, Kawahara Y, Yamashita O, Fukuma R, Yamamoto S, Saitoh Y, et al. Neural decoding of electrocorticographic signals using dynamic mode decomposition. Journal of Neural Engineering. 2020;17(3):036009. doi:10.1088/1741-2552/ab8910.

56. Cohen MX. Comparison of linear spatial filters for identifying oscillatory activity in multichannel data. Journal of Neuroscience Methods. 2017;278:1–12. doi:10.1016/j.jneumeth.2016.12.016.

57. De Schutter B. Minimal state-space realization in linear system theory: an overview. Journal of computational and applied mathematics. 2000;121(1-2):331–354.

58. Golub GH, Loan CFV. Matrix Computations (Johns Hopkins Studies in the Mathematical Sciences). Johns Hopkins University Press; 2012.

59. Qyinn A, Hymers M. SAILS: Spectral Analysis In Linear Systems. Journal of Open Source Software. 2020;5(47):1982. doi:10.21105/joss.01982.

60. Colclough GL, Woolrich MW, Tewarie PK, Brookes MJ, Qyinn AJ, Smith SM. How reliable are MEG resting-state connectivity metrics? NeuroImage. 2016;138:284–293. doi:10.1016/j.neuroimage.2016.05.070.

61. Bürkner PC. brms: An R Package for Bayesian Multilevel Models Using Stan. Journal of Statistical Software. 2017;80(1). doi:10.18637/jss.v080.i01.

62. Bürkner PC. Advanced Bayesian Multilevel Modeling with the R Package brms. The R Journal. 2018;10(1):395. doi:10.32614/rj-2018-017.

63. Vehtari A, Gelman A, Gabry J. Practical Bayesian model evaluation using leave-one-out cross-validation and WAIC. Statistics and Computing. 2017;27(5):1413–1432. doi:10.1007/s11222-016-9696-4.

64. Tzourio-Mazoyer N, Landeau B, Papathanassiou D, Crivello F, Etard O, Delcroix N, et al. Automated Anatomical Labeling of Activations in SPM Using a Macroscopic Anatomical Parcellation of the MNI MRI Single-Subject Brain. NeuroImage. 2002;15(1):273–289. doi:10.1006/nimg.2001.0978.

65. Woolrich M, Hunt A, Barnes G. MEG beamforming using Bayesian PCA for adaptive data covariance matrix regularization. NeuroImage. 2011;57:1466–1479. doi:10.1016/j.neuroimage.2011.04.041.

66. Akaike H. A new look at the statistical model identification. IEEE Transactions on Automatic Control. 1974;19(6):716–723. doi:10.1109/tac.1974.1100705.

67. Brovelli A, Ding M, Ledberg A, Chen Y, Nakamura R, Bressler SL. Beta oscillations in a large-scale sensorimotor cortical network: Directional influences revealed by Granger causality. Proceedings of the National Academy of Sciences. 2004;101(26):9849–9854. doi:10.1073/pnas.0308538101.

68. Schlögl A, Supp G. Analyzing event-related EEG data with multivariate autoregressive parameters. In: Neuper C, Klimesch W, editors. Progress in Brain Research. vol. 159 of Event-Related Dynamics of Brain Oscillations. Elsevier; 2006. p. 135–147. Available from: http://www.sciencedirect.com/science/article/pii/S0079612306590090.

69. Ding M, Bressler SL, Yang W, Liang H. Short-window spectral analysis of cortical event-related potentials by adaptive multivariate autoregressive modeling: data preprocessing, model validation, and variability assessment. Biological Cybernetics. 2000;83(1):35–45. doi:10.1007/s004229900137.

70. Jansen BH. Time series analysis by means of linear modelling. In: Weitknat R, editor. Digital Biosignal Processing (Techniques in the Behavioral and Neural Sciences). Elsevier Science Ltd; 1991. p. 157–180.

71. Lütkepohl H. New Introduction to Multiple Time Series Analysis. Springer; 2007. Available from: https://www.xarg.org/ref/a/3540262393/.

72. Durbin J, Watson GS. Testing for Serial Correlation in Least Squares Regression I. Biometrica. 1950;37:409–428.

73. Nichols TE, Holmes AP. Nonparametric permutation tests for functional neuroimaging: a primer with examples. Hum Brain Mapp. 2002;15(1):1–25.

